# Microbial Biogeography Across the Bovine Body: A Meta-analysis of 27 Anatomical Niches

**DOI:** 10.64898/2026.07.28.741381

**Authors:** Godson Aryee, Devin B. Holman, Carl R. Dahlen, Samat Amat

**Affiliations:** Department of Animal Sciences, North Dakota State University, Fargo, ND, USA; Lacombe Research and Development Centre, Agriculture and Agri-Food Canada, Lacombe, AB, Canada

**Keywords:** Meta-analysis, bovine microbiome, anatomical niches, host-associated microbiota, microbial ecology, 16S rRNA gene amplicon sequencing

## Abstract

Cattle harbor diverse bacterial communities across the gastrointestinal, respiratory, reproductive, mammary and other anatomical systems, but body-wide microbial biogeography remains poorly defined. To address this knowledge gap, we conducted a meta-analysis of publicly available bovine 16S rRNA gene amplicon sequencing data from 5,637 samples from 47 studies across six geographic regions and 27 anatomical sample types. Bacterial community structure differed significantly among sample types (PERMANOVA, R^2^ = 0.245, *P* = 0.0001), indicating spatial organization of bacterial communities across the bovine body, although study-level effects also contributed substantially to community variation. Communities were generally more similar within than between anatomical systems. Bacterial richness, diversity, taxonomic composition, and indicator taxa varied among sample types, with gastrointestinal, mammary-associated, ocular, and hoof microbiota exhibiting greater diversity than microbiota from liver, joint, and several reproductive samples. Distinct bacterial communities characterized the gastrointestinal, respiratory, reproductive, and mammary systems, as well as other anatomical sites. Despite these differences, several bacterial taxa were shared across multiple anatomical niches, particularly among male and female reproductive sites and among mammary-associated niches. This study provides a comprehensive body-wide characterization of bacterial biogeography in cattle and establishes a baseline for future studies of bovine microbial ecology and host–microbiome interactions.

## INTRODUCTION

There are approximately 1.6 billion cattle worldwide, making them one of the most important livestock species for global food production [1]. Dairy and beef cattle contribute substantially to human nutrition and food security [2, 3]. Yet the cattle industry faces mounting pressures related to antimicrobial resistance, environmental sustainability, animal health, and the growing demand for animal protein [4]. Traditionally, improvements in cattle productivity have been achieved through genetic selection and management practices [5, 6]. However, advances in high-throughput sequencing have revealed that host-associated microbiota also play fundamental roles in host physiology, immunity, reproduction, and health [7–11]. As a result, the bovine microbiome has become an important focus of research aimed at improving cattle health, productivity, and environmental sustainability.

High-throughput sequencing studies have demonstrated that cattle harbor distinct microbial communities throughout the gastrointestinal [8, 9], respiratory [12], reproductive [11], and mammary systems [7]. Microbial communities have also been characterized in samples from other anatomical sites, including the oral cavity [13], ocular surface [14, 15], hoof [16, 17], liver [18, 19], blood [20, 21], and joints [22, 23]. Among these, the rumen microbiome has been the most extensively investigated because of its central role in feed digestion, nutrient metabolism, feed efficiency, and enteric methane emissions [9, 24]. Likewise, respiratory microbiota have been widely studied because of their associations with bovine respiratory disease (BRD) [12]. Reproductive tract microbiota have been associated with fertility and reproductive health in both females and males [11, 25–28]. Mammary-associated microbiota have also been linked to milk production traits and udder health [7, 29]. In addition, microbial communities inhabiting other body sites are increasingly recognized as potential contributors to tissue homeostasis and disease resistance [17, 30, 31].

Despite rapid advances in bovine microbiome research, most studies have focused on individual anatomical sites or single organ systems. Consequently, relatively little is known about how bacterial communities are organized across the bovine body, whether common ecological patterns exist among anatomically distinct habitats, or which bacterial taxa are consistently shared across body systems. Defining microbial biogeography across the bovine body is therefore important because it provides a baseline for comparing bacterial communities among anatomical niches and identifying site-specific and shared bacterial taxa.

Comparative analyses across multiple anatomical niches may also improve our understanding of potential microbial connectivity within the host. Emerging evidence from both human and animal studies indicates that gastrointestinal microbial communities may influence distant organs through gut–organ axes, including the gut–brain, gut–lung, gut–liver, gut– mammary, and gut–reproductive axes [32–36] (Fig. 1). Although many gut-organ interactions are thought to be mediated through microbial metabolites, immune signaling, and host physiological responses, recent studies have also reported taxonomic overlap among the gastrointestinal, respiratory, reproductive, and other body sites in cattle [37–39], indicating that some bacterial taxa may occur across multiple anatomical niches. However, detecting the same bacterial taxonomic groups at multiple sites does not establish that the same strains are present, nor does it demonstrate microbial transmission or functional connectivity among anatomical sites.

**Figure 1.**
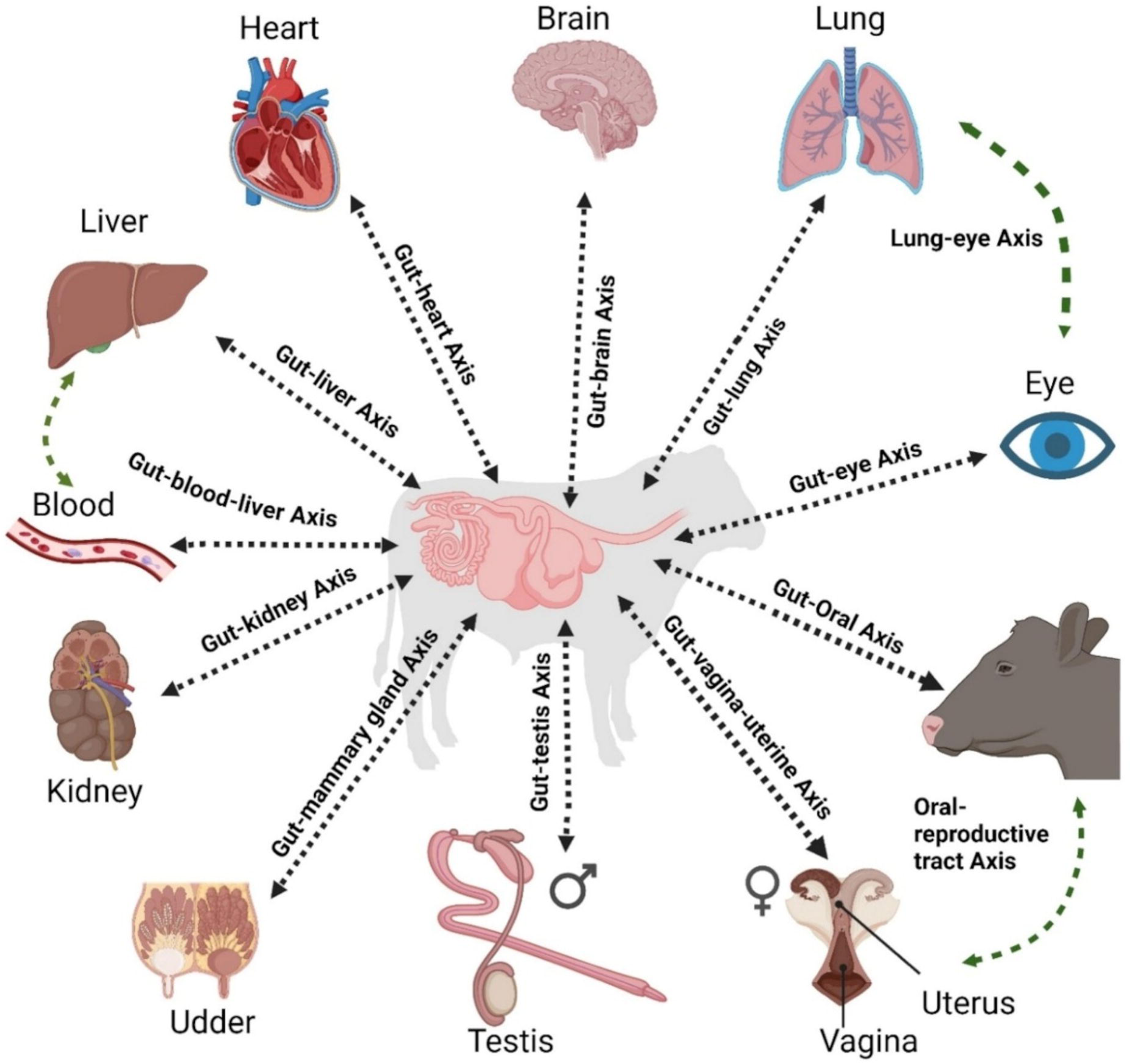
A schematic overview of the gut-all other organs-microbiome axes. This figure shows the gut–organ axis in cattle, showing how the gut communicates with the brain, lungs, liver, heart, kidney, mammary gland, reproductive tract, eye, and blood through interconnected pathways collectively referred to as gut–organ axes (e.g., gut–brain, gut–lung, gut–liver, gut– mammary, gut–reproductive, and gut–blood axes). The diagram emphasizes the concept that microbial communities in the gut may play a key role in whole-body regulation and developmental programming in cattle.

Beyond their potential role in inter-organ communication, maternal microbiota are increasingly recognized as contributors to offspring developmental programming and microbiome establishment, whereas evidence for contributions from paternal microbiota is emerging [40–43]. Characterizing microbial communities across multiple parental anatomical niches may therefore help identify anatomical sites that serve as microbial sources for the developing offspring microbiome. Moreover, bacterial taxa detected across multiple body sites may exhibit different ecological roles depending on the anatomical niche they occupy, functioning as commensals in one habitat while being associated with disease under different ecological or host conditions [44]. A body-wide view of microbial distribution therefore provides an important baseline for understanding host-associated microbial ecology and for generating hypotheses regarding microbial transmission, niche adaptation, and host–microbiome interactions.

To address this knowledge gap, we conducted a meta-analysis of publicly available bovine 16S rRNA gene amplicon sequencing datasets reported in studies published from 2020 through 2024. The combined dataset comprised 5,637 samples from 47 studies representing 27 anatomical niches in beef and dairy cattle across six geographic regions. Specifically, we aimed to determine whether bacterial communities are anatomically structured across the bovine body, identify bacterial taxa associated with individual sample types, and quantify taxonomic overlap among related anatomical niches. This study provides a comprehensive characterization of bacterial biogeography across the bovine body and establishes a framework for future studies of the bovine microbiome.

## MATERIALS AND METHODS

### Acquisition of 16S rRNA gene amplicon sequencing datasets

To assemble a large-scale dataset of bovine microbiota across anatomical niches, we searched public sequence repositories and literature databases, including the National Center for Biotechnology Information Sequence Read Archive (NCBI SRA), the European Nucleotide Archive (ENA), PubMed, and Google Scholar. Searches were performed using combinations of the following keywords and phrases: "16S rRNA sequencing," "bovine," "gut microbiome/microbiota," "respiratory microbiome/microbiota," "ocular microbiome/microbiota," "reproductive microbiome/microbiota," "udder microbiota," "mammary gland microbiota," "milk microbiota," "liver microbiota," "blood microbiota," "hoof microbiota," "joint microbiota," "beef," "dairy," and "dual-purpose." This search identified 100 studies published from January 2020 to April 2024, inclusive, for initial screening (Table S1). Studies published prior to 2020 were excluded to focus on recent 16S rRNA gene amplicon sequencing datasets and to avoid overlap with our previous meta-analysis of the bovine gastrointestinal tract [45]. To be included in this meta-analysis, studies were required to: (i) use high-throughput sequencing of 16S rRNA gene amplicons, (ii) have paired-end 16S rRNA sequence data, (iii) be conducted on domestic cattle (*Bos taurus*), including beef, dairy, or dual-purpose animals, (iv) provide clearly described associated metadata, and (v) have sequence data publicly available in the SRA or ENA. Of the 100 studies initially identified, 47 met the inclusion criteria and were included in the final analysis (Table S1). Sequence datasets were downloaded from the SRA using the SRA Toolkit or directly from the ENA, depending on where they were originally deposited.

### Processing and analysis of 16S rRNA gene amplicon sequencing data

All 16S rRNA gene sequencing datasets were processed using QIIME 2 v.2024.10 [46] following Dong et al. [47]. Briefly, paired-end reads were merged using the VSEARCH join-pairs method implemented in QIIME 2. Merged reads were then quality filtered using a Phred quality score threshold of 25 and dereplicated prior to operational taxonomic unit (OTU) clustering. Dereplicated sequences were clustered into OTUs at 97% sequence identity using closed-reference OTU picking implemented VSEARCH in QIIME 2 [48] against the SILVA SSU rRNA database release 138.2 [49]. Chimeric sequences were identified using de novo UCHIME implemented in QIIME 2 VSEARCH and removed. A closed-reference OTU-picking strategy was employed to assign sequences generated from different 16S rRNA gene hypervariable regions to a common reference-based OTU framework.

Mapping sequences to a single curated reference database at 97% sequence identity enabled comparisons across studies, although reads that did not match a reference sequence at the selected threshold were excluded [50–52]. OTUs classified as chloroplast or mitochondria were subsequently removed. Archaeal OTUs were also excluded because the primer sets used across the included studies did not consistently capture archaeal taxa. After samples containing fewer than 1,000 sequences were removed, 5,637 samples were retained for taxonomy-based analyses. For alpha- and beta-diversity analyses, samples were rarefied to 5,000 sequences per sample, resulting in a dataset of 5,237 samples. Relative abundance, heat map, and indicator-taxon analyses were performed using the unrarefied OTU table after samples containing fewer than 1,000 sequences were removed.

Alpha-diversity metrics, including observed OTUs, Shannon diversity, and inverse Simpson diversity, were calculated from the rarefied OTU table in R v.4.5.2 using the vegan package v.2.7.3. To visualize sample type-associated taxonomic patterns within each body system, the unrarefied OTU table, after removal of samples containing fewer than 1,000 sequences, was transformed to within-sample relative abundance. Mean relative abundance was then calculated for each OTU within each sample type. For each OTU, the mean relative abundance was transformed as log_10_(mean relative abundance + 1 x 10^-^□). OTUs were ranked according to the variance of these transformed values across sample types, and the 50 most variable OTUs within each body system were selected for visualization. For each selected OTU, the log10-transformed mean relative abundance values were standardized across sample types as row-wise z-scores to visualize relative differences among sample types within each body system.

### Statistical analysis

Bray–Curtis dissimilarities were calculated from the rarefied OTU table using phyloseq v. 1.50.0 [53] and bacterial community structure was visualized using non-metric multidimensional scaling (NMDS) implemented in vegan. Permutational multivariate analysis of variance (PERMANOVA) based on Bray–Curtis dissimilarities was performed using the adonis2 function in vegan to assess whether bacterial community structure differed among sample types within each body system and among anatomical niches across the bovine body. Statistical significance was assessed using 9,999 permutations. For the whole-body analysis, sample type and study were included as sequential terms in the PERMANOVA models. Models were fitted with sample type entered before study and, separately, with study entered before sample type to assess the sensitivity of explained variation to term order.

Genus-level indicator analysis was performed in R using the indicspecies package v.1.8.0 [54]. The analysis was conducted using genus-level relative abundances after retaining genera detected in at least five samples overall and with a prevalence of at least 10% in one or more sample types. Sample type-specific indicator genera were then identified using the group-equalized indicator value statistic (IndVal.g), with associations restricted to individual sample types rather than combinations of sample types. Statistical significance was assessed using 9,999 permutations. The resulting P values were adjusted for multiple testing using the Benjamini-Hochberg procedure.

## RESULTS

### Dataset composition and sample distribution

A total of 47 studies met the inclusion criteria. After removal of samples with fewer than 1,000 sequences, the final dataset comprised 5,637 samples collected from cattle in 13 countries spanning six geographic regions: Africa, Asia, Europe, North America, Oceania, and South America (Fig. 2; Table S1). Of these, 5,237 samples had at least 5,000 sequences and were retained for rarefied alpha- and beta-diversity analyses. The final dataset had a mean sequencing depth of 53,619 ± 906 reads per sample (mean ± SEM). More than half of the samples originated from the United States (52.5%; n = 2,958), followed by Canada (14.4%; n = 810). The datasets targeted the V1–V3 (0.5%; n = 26), V3 (0.3%; n = 15), V3–V4 (30.6%; n = 1,725), or V4 (68.7%; n = 3,871) hypervariable regions of the 16S rRNA gene (Table S1).

**Figure 2.**
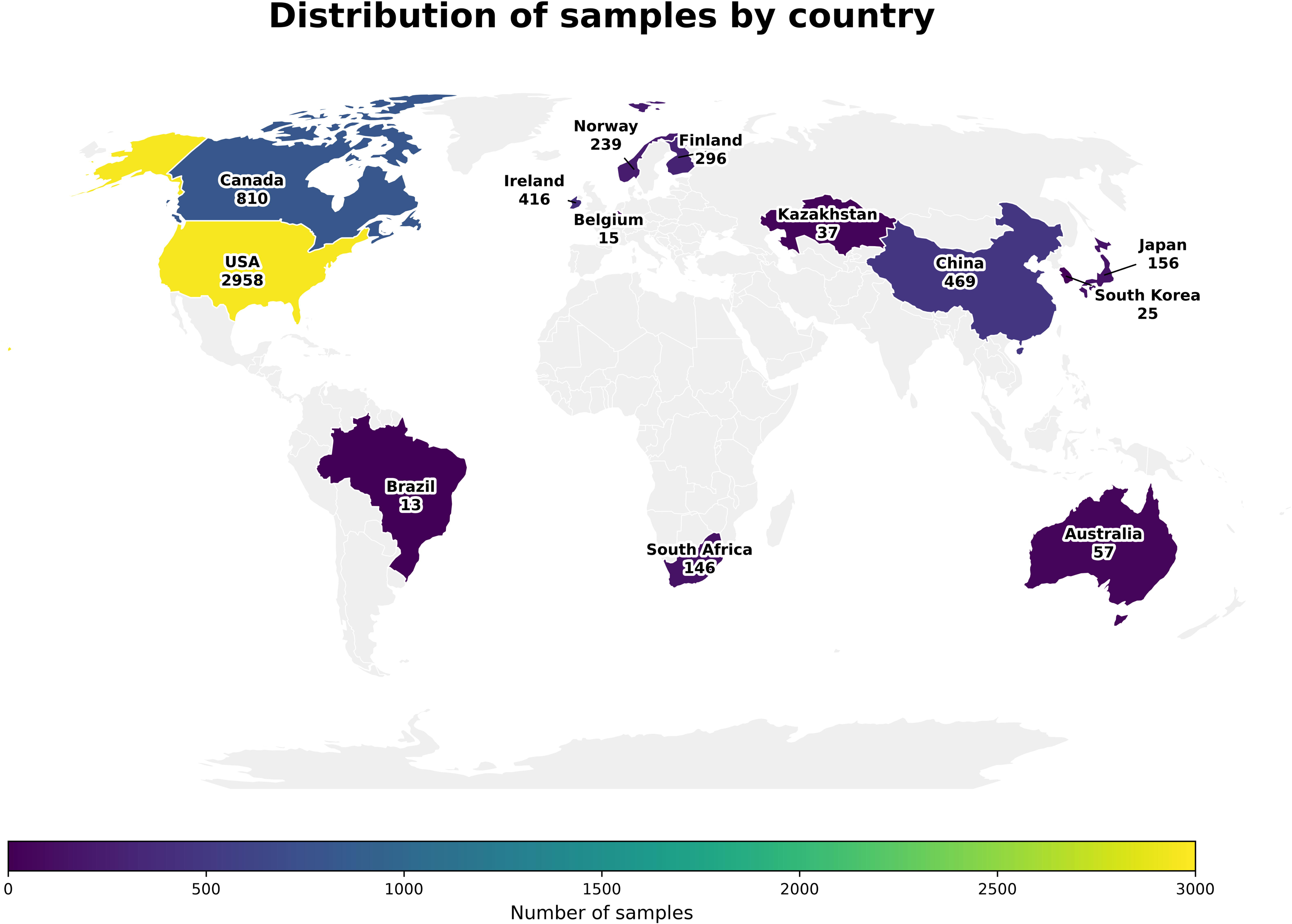
Geographic distribution of samples included in the meta-analysis, with countries shaded according to the number of samples contributed. Additional dataset details are provided in Table S1.

The compiled dataset represented 27 anatomical niches spanning the gastrointestinal tract (38.0%; n = 2,142), ocular surface (25.6%; n = 1,444), respiratory tract (13.8%; n = 776), reproductive tract (12.5%; n = 705), mammary-associated sites (4.4%; n = 249), and other body sites (5.7%; n = 321), including hoof, liver, blood, and joint samples (Table S1). Samples were obtained primarily from beef cattle (65.9%; n = 3,717), followed by dairy cattle (23.4%; n = 1,319). Cattle type was not specified for 10.5% of samples (n = 594) and dual-purpose cattle accounted for 0.1% of samples (n = 7).

### Bacterial biogeography across 27 bovine anatomical niches

When sample type was entered first in the sequential PERMANOVA model, it explained 24.5% of the variation in bacterial community structure (PERMANOVA, R^2^ = 0.245, *P* = 0.0001; Fig. 3). Bacterial communities clustered primarily according to anatomical origin, with samples generally grouping by gastrointestinal, respiratory, reproductive, mammary-associated, and other anatomical categories. Overlap was observed among several anatomically related niches, particularly within the reproductive and mammary-associated sites, whereas other niches remained highly distinct. For example, ocular surface and endometrial samples displayed distinct clustering, consistent with anatomical structuring of bacterial communities across the bovine body. However, because several sample types were represented by only one or a small number of studies, study-level effects were also substantial. When study was entered first in the model, it explained 40.0% of the variation (R^2^ = 0.400), while sample type explained an additional 3.6% of variation (R^2^ = 0.036; *P* = 0.0001).

**Figure 3.**
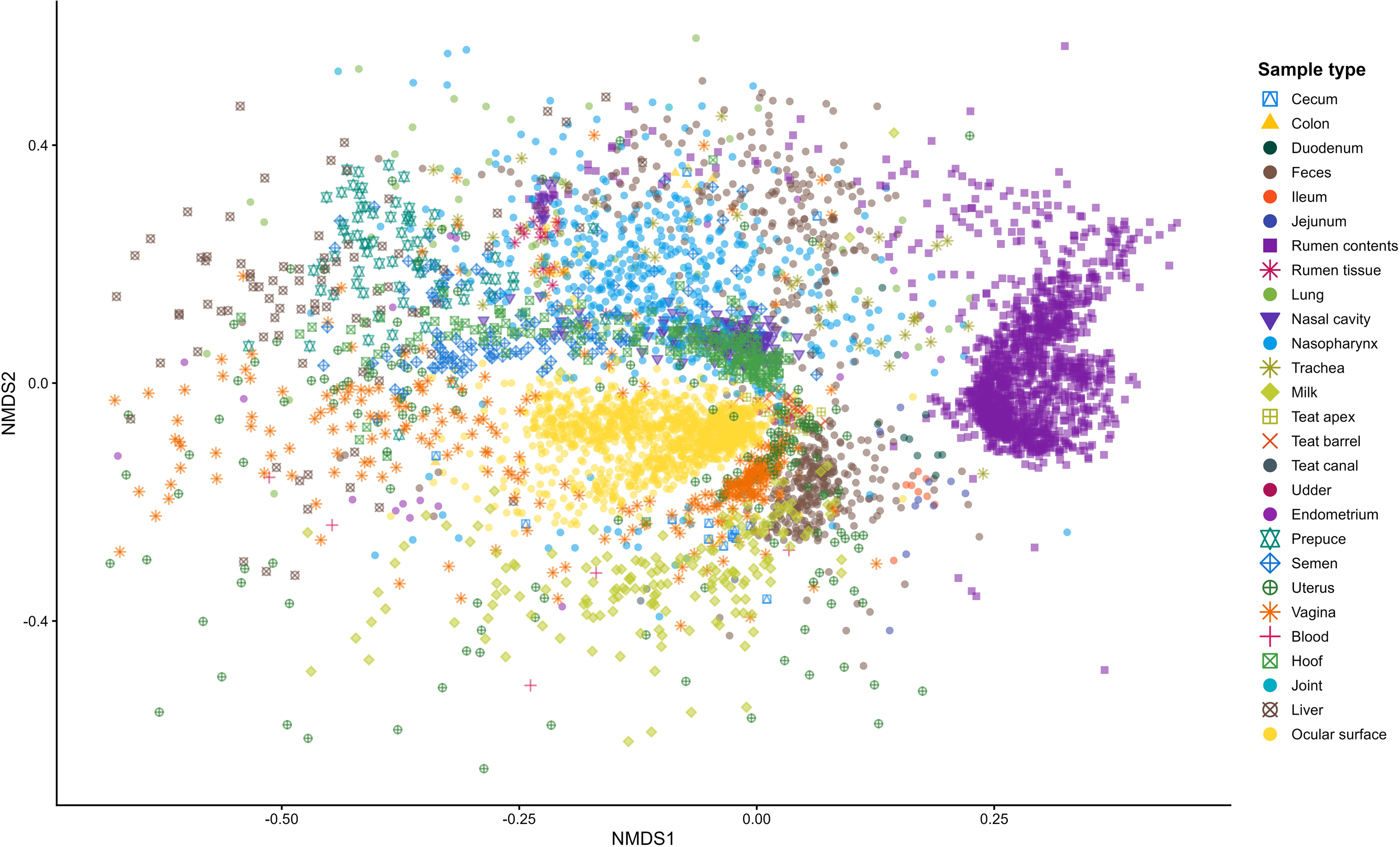
Non-metric multidimensional scaling (NMDS) ordination based on Bray-Curtis dissimilarities in bacterial community composition across 27 bovine anatomical sites (n = 5,237).

Microbial richness and diversity also varied substantially among anatomical niches (Fig. 4). Gastrointestinal, mammary-associated, ocular, and hoof microbiota displayed the greatest richness and diversity, whereas liver, blood, joint, and several reproductive sample types harbored comparatively lower diversity. Within the gastrointestinal tract, bacterial diversity was greatest in the rumen and hindgut and lower in the small intestine. Outside the gastrointestinal tract, vaginal, uterine, teat-associated, and ocular microbiota also showed relatively high diversity, whereas semen, preputial, liver, blood, and joint microbiota were comparatively less diverse.

**Figure 4.**
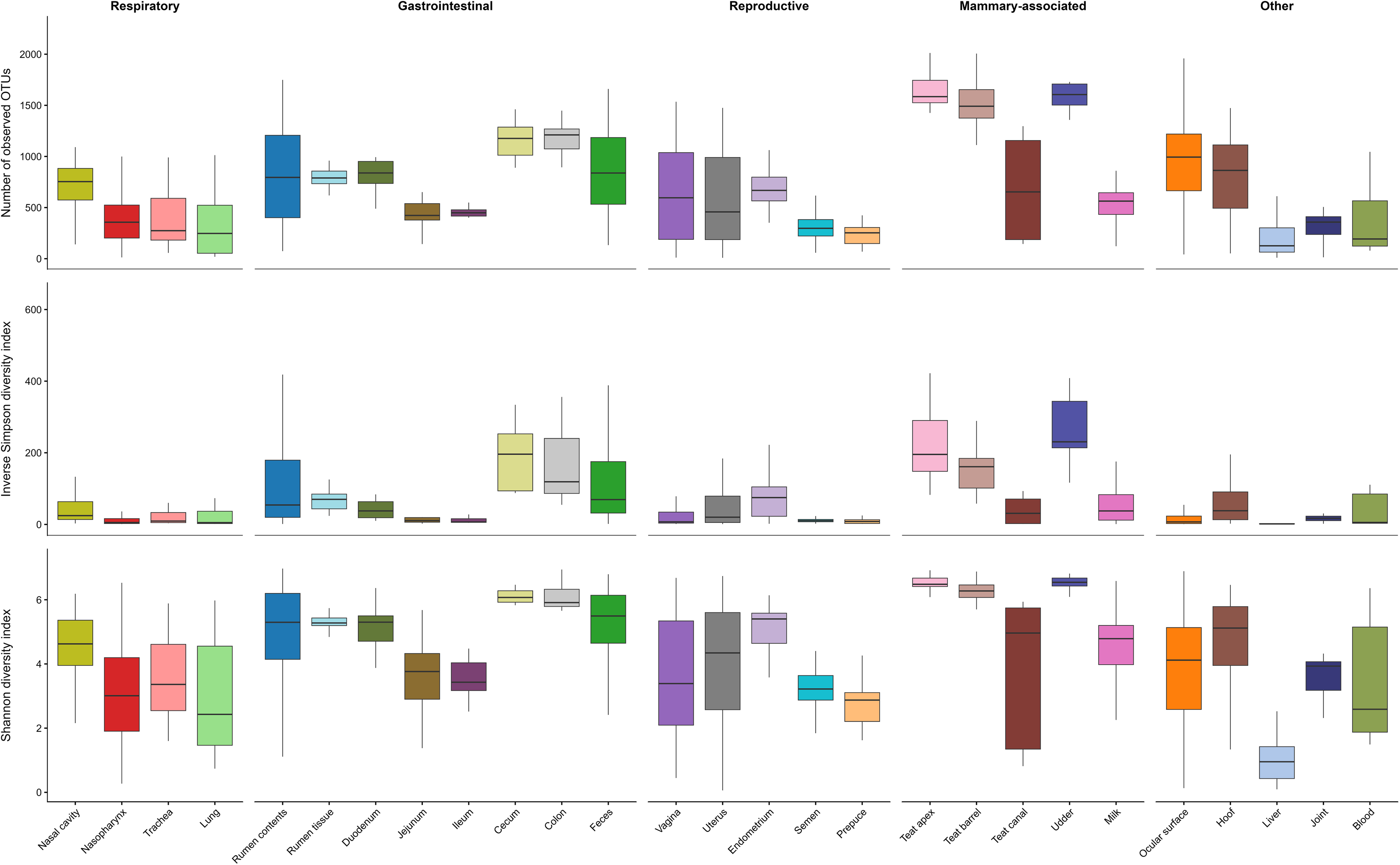
Comparison of alpha diversity measures across 27 bovine anatomical sites (n = 5,237). Bacterial richness was assessed using the number of observed OTUs, and diversity was assessed using the inverse Simpson and Shannon diversity indices. Boxplots show the distribution of each metric across sample types grouped by body system or anatomical category. Boxes represent the interquartile range, horizontal lines indicate medians, and whiskers extend to 1.5 times the interquartile range.

Despite significant differences in community structure and considerable variation in diversity, bacterial communities across the bovine body were dominated by three major phyla: Bacillota, Pseudomonadota, and Bacteroidota, although their relative abundances varied considerably among anatomical niches (Fig. 5). Gastrointestinal microbiota were characterized by higher relative abundances of Bacillota and Bacteroidota, whereas Pseudomonadota predominated in several respiratory, reproductive, and blood sample types. Mammary-associated, ocular, hoof, liver, and joint microbiota also displayed distinct phylum-level profiles.

**Figure 5.**
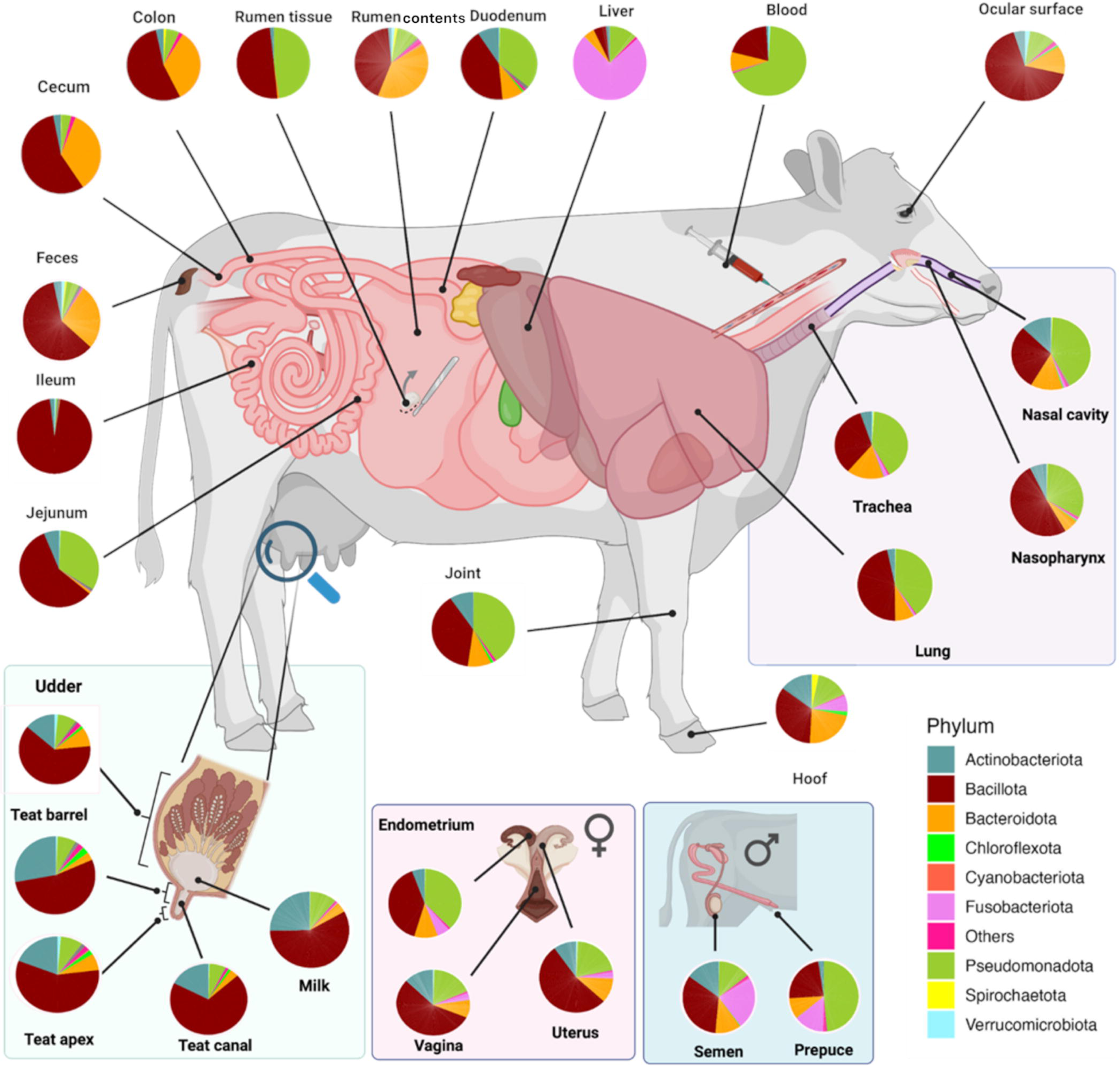
Phylum-level composition of bacterial communities across 27 bovine anatomical sites (n = 5,637). Pie charts show the relative abundance of the dominant bacterial phyla within each anatomical site. Phyla are indicated by color, and lower-abundance phyla are grouped as “Others.”

No individual OTU was detected in more than 63% of all samples. The most globally prevalent OTU was classified as *Mogibacterium* and was detected in 62.8% of samples overall. However, several other OTUs were detected across all 27 anatomical sample types (Table S2). Among these, the most consistently distributed OTU was affiliated with the [*Eubacterium*] *tenue* group, which was present at ≥5% prevalence in all sample types and detected in 58.8% of samples overall. Additional OTUs classified as *Turicibacter*, *Romboutsia*, *Bifidobacterium*, *Mailhella*, and *Streptococcus* were also detected across all 27 sample types, although their prevalence varied among anatomical niches.

The distribution of selected bacterial genera that include recognized bovine pathogens differed substantially among anatomical niches (Fig. 6). *Fusobacterium* and *Mycoplasma* were the most broadly distributed genera, occurring across multiple body systems, whereas *Mannheimia, Pasteurella, Moraxella, Ureaplasma, Trueperella,* and *Histophilus* exhibited more anatomically restricted distributions. For example, *Fusobacterium* was most abundant in liver samples but was also detected in reproductive and hoof microbiota. *Mycoplasma* was most abundant in respiratory and ocular niches. *Ureaplasma* was largely restricted to the female reproductive tract, and *Mannheimia* and *Pasteurella* were primarily associated with the upper respiratory tract.

**Figure 6.**
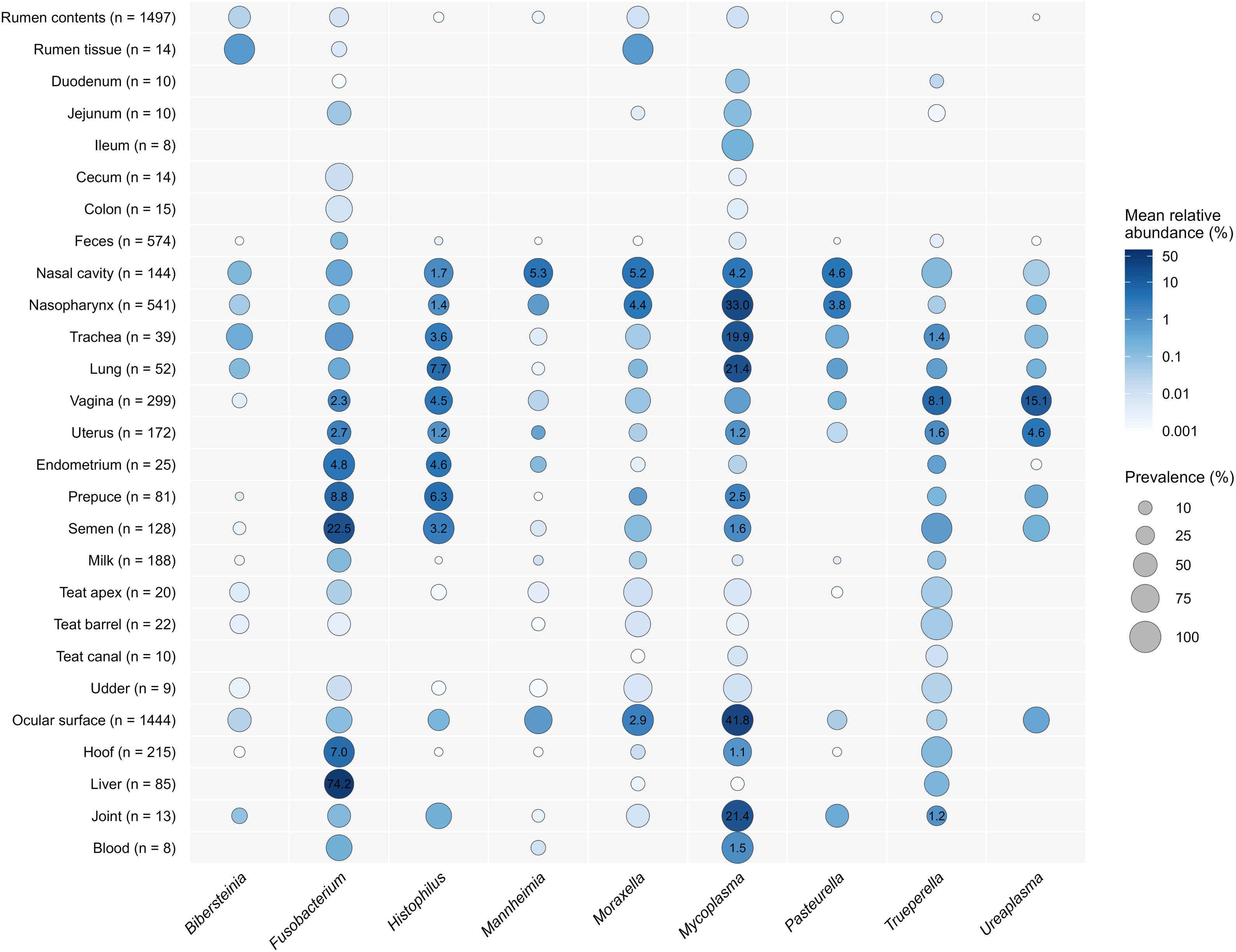
Relative abundance and prevalence of selected pathogen-associated bacterial genera across bovine anatomical sample types. Bubble colour indicates mean relative abundance (%) within each sample type and bubble size indicates prevalence, defined as the percentage of samples in which the genus was detected at ≥0.001% relative abundance. Numeric labels indicate mean relative abundance (%) and are shown only for genus–sample type combinations with a mean relative abundance >1%.

A total of 844 significant indicator genera were identified across the 27 anatomical sample types examined, with the largest numbers identified in gastrointestinal tract and mammary-associated sample types, followed by respiratory, reproductive, and other anatomical sites. (Fig. 7; Table S4). Among sample types represented by more than 100 samples, notable indicators included *Prevotella* in rumen contents*, Fournierella* in feces, *Faucicola* in the nasal cavity, *Trueperella* and *Ureaplasma* in the vagina, S5-A14a (family *Anaerovoracaceae*) in semen, *Lactococcus* and *Weissella* in milk, *Mycoplasma* on the ocular surface, and *Ezakiella* on the hoof.

**Figure 7.**
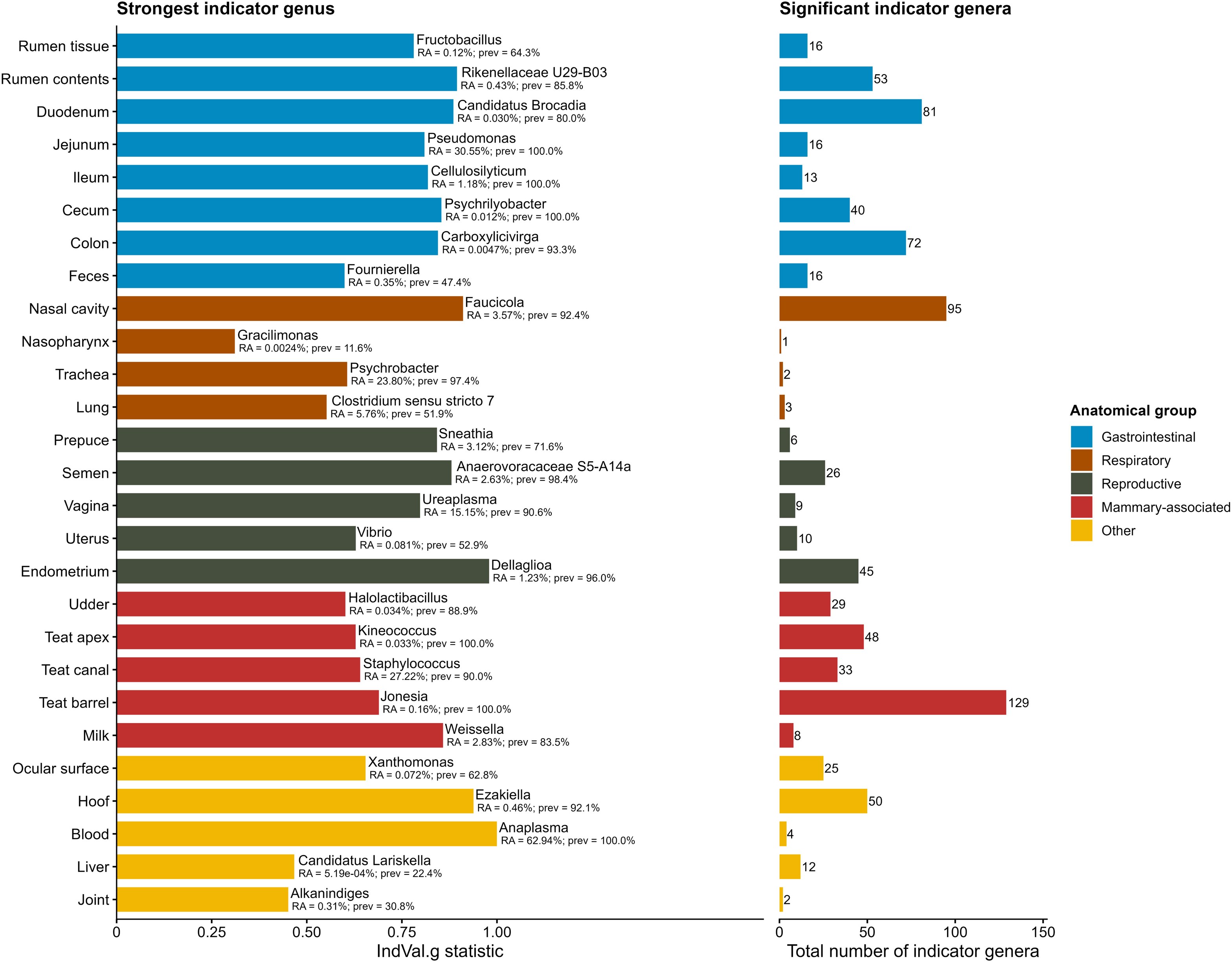
Indicator genera associated with bovine anatomical sample types (n = 5,637). Indicator analysis was performed at the genus level using group-equalized indicator values (IndVal.g; FDR < 0.05). For each sample type, the genus with the highest IndVal.g statistic is shown, along with its mean relative abundance and prevalence within that sample type. The right-hand bar shows the total number of significant indicator genera identified for each sample type. Bar colour indicates anatomical group. RA, relative abundance; prev, prevalence.

These sample type-associated genera further supported strong anatomical niche specialization of the bovine microbiota. Together, the observed differences in bacterial community composition, diversity, taxonomic profiles, and indicator genera revealed clear spatial organization of bacterial communities across the bovine body. We next examined microbial organization within the respiratory, gastrointestinal, reproductive, mammary-associated, and other anatomical systems to characterize finer-scale variation among individual niches.

### Bacterial community variation within major bovine body systems

#### Bacterial biogeography of the respiratory tract

Bacterial community composition differed significantly among airway sites (PERMANOVA, R^2^ = 0.11, *P* = 0.0001; Fig. S1A). The nasal cavity was characterized by a distinct bacterial community, whereas nasopharyngeal, tracheal, and lung microbiota showed greater overlap. Bacterial richness and diversity progressively declined from the upper to the lower respiratory tract, with the nasal cavity exhibiting the greatest bacterial diversity (Fig. 4). Respiratory microbiota were dominated by Bacillota and Pseudomonadota across all airway compartments (Fig. S1B). At the genus level, *Mycoplasma* predominated throughout the respiratory tract, while the nasal cavity harbored a more diverse bacterial community that included *Acinetobacter, Corynebacterium*, *Moraxella*, *Pasteurella,* and *Streptococcus* (Fig. S1C; Table S3). In contrast, tracheal and lung microbiota were characterized by high relative abundances of *Mycoplasma*, *Psychrobacter*, and *Streptococcus*. At the OTU level, *Moraxella*-, *Pasteurella*-, and *Mannheimia*-associated OTUs showed higher relative abundances in the upper respiratory tract, while *Burkholderia-Caballeronia-Paraburkholderia*-associated OTUs showed higher relative abundances in lung samples (Fig. S2).

#### Bacterial biogeography within the gastrointestinal tract

Bacterial community composition also differed significantly among sites within the gastrointestinal tract (PERMANOVA, R^2^ = 0.09, *P* = 0.0001; Fig. 8A). Microbiota from rumen contents formed the most distinct cluster, while small intestinal microbiota clustered closely together. Cecal and colonic microbiota occupied positions intermediate between the small intestine and feces, indicating gradual changes in bacterial community composition along the gastrointestinal tract. Bacterial richness and diversity also varied among gastrointestinal sites (Fig. 4). Richness was greatest in rumen contents, cecal, colonic, and fecal microbiota, whereas diversity was generally greatest in cecal and colonic microbiota and lower in the small intestine. Bacillota, Pseudomonadota, and Bacteroidota dominated bacterial communities throughout the gastrointestinal tract, although their relative abundances varied among gastrointestinal sites (Fig. 8B). The relative abundance of Pseudomonadota was higher in rumen tissue and the duodenum than at most other gastrointestinal sites. Bacillota and Bacteroidota predominated across most other gastrointestinal sites. At the genus level, *Prevotella* predominated in rumen contents, while hindgut microbiota contained higher relative abundances of *Bacteroides*, *Rikenellaceae* RC9 gut group, *Christensenellaceae* R-7 group, and other obligate anaerobic taxa. Small intestinal microbiota showed lower relative abundances of the dominant rumen and hindgut-associated genera (Fig. 8C; Table S4).

**Figure 8.**
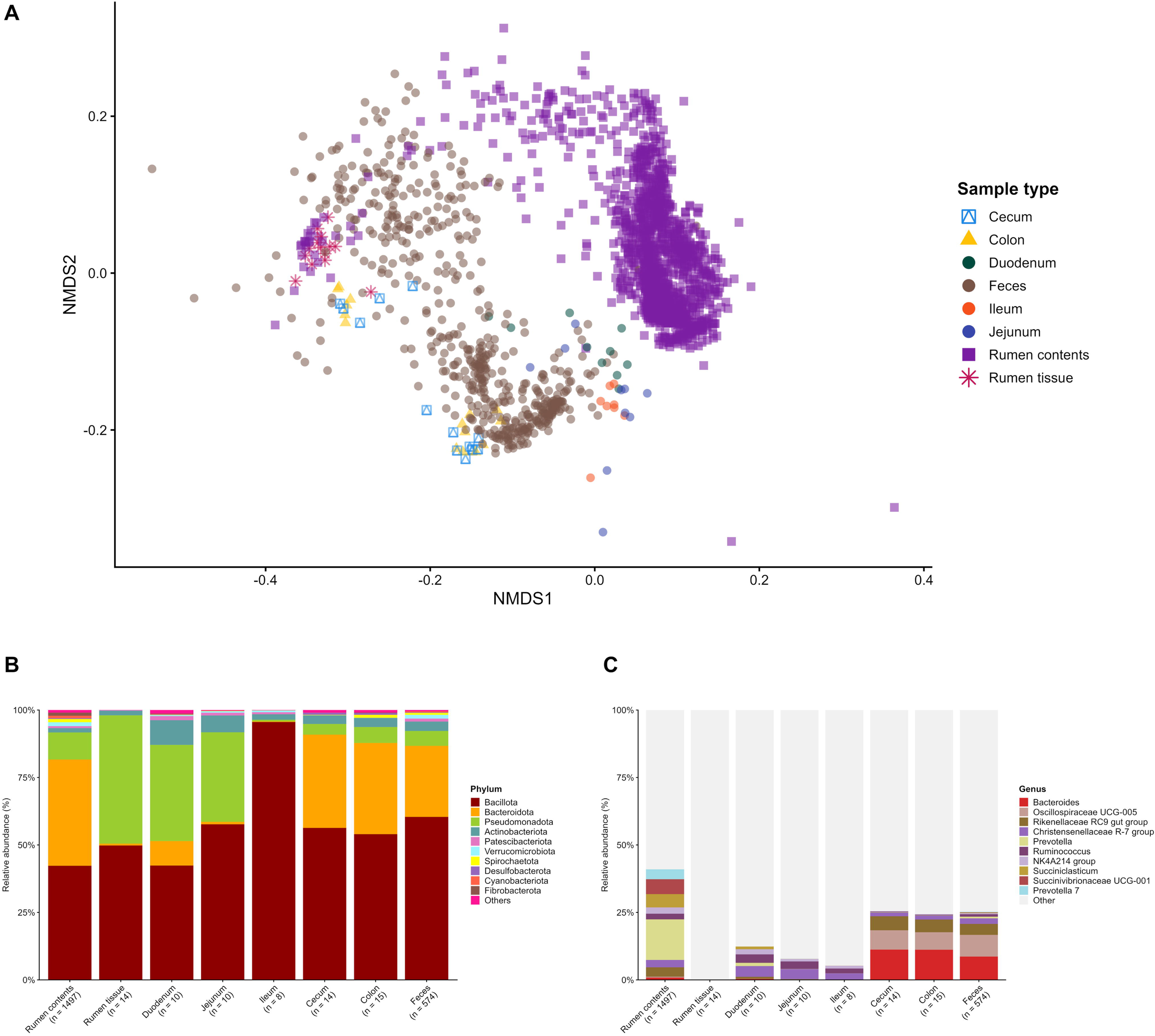
Bacterial microbiota associated with bovine gastrointestinal tract sample types. **A)** Non-metric multidimensional scaling (NMDS) ordination based on Bray-Curtis dissimilarities in bacterial community structure among gastrointestinal tract samples (n = 2,062). **B)** Phylum-level relative abundance (%) by sample type. **C**) Genus-level relative abundance (%) by sample type. For stacked bar plots (n = 2,142), the most abundant taxa are shown individually and remaining taxa are grouped as “Other” or “Others.”

OTU-level abundance patterns also differed along the gastrointestinal tract (Fig. S3). Rumen contents microbiota had OTU profiles distinct from those of the small intestine. Cecal, colonic, and fecal microbiota shared similar abundance patterns, with higher relative abundances of OTUs associated with *Rikenellaceae* RC9 gut group, *Alistipes*, *Prevotellaceae* UCG-004, and other obligate anaerobes. Although fecal microbiota shared many abundant OTUs with the hindgut, they also contained several distinct OTUs, consistent with continued spatial differentiation along the distal gastrointestinal tract.

#### Bacterial biogeography within the reproductive tract

Bacterial community structure differed significantly among male and female reproductive sites (PERMANOVA, R^2^ = 0.165, *P* = 0.0001; Fig. S4A). Semen and preputial samples clustered closely together, while vaginal and uterine samples partially overlapped. Endometrial samples remained distinct from the other reproductive sites. Bacterial richness and diversity also varied among reproductive sites (Fig. 4). Vaginal, uterine, and endometrial samples displayed greater richness and diversity than semen and preputial samples.

Bacillota, Pseudomonadota, and Bacteroidota were the dominant phyla in the reproductive microbiota, although their relative abundances differed among reproductive sites (Fig. S4B). The relative abundance of Bacillota was higher in female reproductive sites, whereas Pseudomonadota represented a larger proportion of preputial microbiota. At the genus level, vaginal and uterine microbiota included *Ureaplasma*, *Histophilus*, and *Helcococcus*. Endometrial microbiota included *Streptococcus* and *Escherichia-Shigella*. Semen and preputial microbiota were characterized by *Fusobacterium* and *Escherichia-Shigella*, respectively. Several bacterial genera were shared across male and female reproductive sites (Fig. S4C; Table S4).

Defining an OTU as shared when it was detected in at least one sample from each of the two sample types being compared, vaginal and uterine microbiota shared the greatest number of OTUs (14,450), followed by vaginal and semen microbiota (9,023) (Fig. S5). A total of 1,709 OTUs were shared among the vagina, uterus, semen, and prepuce. OTU-level abundance patterns differed across reproductive sites but also showed considerable similarity among anatomically related sites (Fig. S6). Vaginal and uterine microbiota included OTUs affiliated with *Streptococcus*, *Bifidobacteriaceae*, *Corynebacteriaceae*, *Prevotella*, and *Escherichia-Shigella*. Semen and preputial microbiota included OTUs associated with *Actinomycetaceae*, *Fusobacteriaceae*, *Porphyromonas*, *Acholeplasma*, and *Mycoplasma*. Endometrial microbiota exhibited a distinct OTU profile that included *Fusobacteriaceae-*, *Bacteroides-*, and *Streptococcus*-associated OTUs. Although each reproductive site displayed distinct bacterial abundance patterns, considerable taxonomic overlap was evident among male and female reproductive sites.

#### Bacterial biogeography within mammary-associated sites

Bacterial community composition differed significantly among mammary-associated sites (PERMANOVA, R^2^ = 0.158, *P* = 0.0001; Fig. S7A). Teat apex, teat barrel, teat canal, and udder skin microbiota clustered closely together, while milk samples formed a relatively distinct and dispersed cluster. Bacterial richness and diversity also varied among mammary-associated sites (Fig. 4). Teat apex, teat barrel, and udder skin microbiota displayed greater richness and diversity than milk, whereas teat canal samples showed high variability in alpha diversity.

Bacillota comprised the largest proportion of mammary-associated microbiota, followed by Actinobacteriota, Pseudomonadota, and Bacteroidota (Fig. S7B). Teat-associated and udder skin microbiota shared several prominent genera, including *Staphylococcus*, *Corynebacterium*, *Acinetobacter*, and *Romboutsia*. Milk microbiota were characterized by *Staphylococcus, Corynebacterium, Bacillus, Lactococcus, Cutibacterium,* and *Weissella* (Fig. S7C; Table S4). OTU-level abundance patterns varied among mammary-associated sites (Fig. S8). Milk microbiota exhibited an OTU profile distinct from those of teat-associated and udder skin microbiota, with several milk-associated OTUs affiliated with *Lawsonella*, *Acinetobacter*, *Corynebacteriaceae*, *Bacillus*, *Weissella*, *Staphylococcus*, and *Lactococcus*. In contrast, teat-associated and udder skin microbiota shared similar OTU profiles that included OTUs affiliated with *Carnobacteriaceae*, *Streptococcus*, *Acinetobacter*, *Corynebacteriaceae*, *Romboutsia*, and *Turicibacter*.

#### Bacterial biogeography across other bovine anatomical sites

For descriptive presentation, ocular surface, hoof, liver, blood, and joint samples, which did not belong to the four multi-site anatomical systems, were grouped as other anatomical sites. Bacterial richness and diversity varied among these anatomical sites (Fig. 4). Ocular and hoof microbiota displayed relatively high bacterial richness and diversity, while liver, blood, and joint samples were comparatively less diverse.

Taxonomic composition also varied among these anatomical sites (Fig. S9A). Bacillota comprised the largest proportion of ocular microbiota. Pseudomonadota was the dominant phylum in blood and joint samples, while liver samples were dominated by Fusobacteriota. Hoof microbiota contained a more even distribution of Bacillota, Pseudomonadota, Bacteroidota, and Actinobacteriota (Fig. S9B). At the genus level, ocular microbiota were dominated by *Mycoplasma*, liver samples were dominated by *Fusobacterium*, and joint microbiota were characterized by *Mycoplasma* and *Psychrobacter* (Fig. S9C; Table S4). OTU-level abundance patterns also differed among these anatomical sites (Fig. S10). Liver samples were characterized by *Fusobacterium*-associated OTUs, ocular microbiota by *Mycoplasma*- and *Moraxella*- associated OTUs, and hoof and joint samples by *Psychrobacter*-associated OTUs.

## Discussion

In this meta-analysis, we integrated publicly available 16S rRNA gene sequencing datasets from 5,637 samples across 47 studies to characterize bacterial biogeography across 27 anatomical niches in cattle. Comparisons among gastrointestinal, respiratory, reproductive, mammary-associated, and other anatomical sites revealed two overarching patterns: anatomical structuring of bacterial communities and the sharing of bacterial taxa among related anatomical niches.

### Whole-body bacterial biogeography and anatomical structuring

The principal finding of this meta-analysis was that bacterial communities differed among anatomical niches across the bovine body. Compartment-specific microbiota have previously been described within individual body systems, including the gastrointestinal tract, respiratory tract, reproductive tract, and mammary gland [7, 8, 11, 55], but direct comparisons across body systems have been limited. By integrating datasets across 27 anatomical niches, this study provides a body-wide comparison of bacterial community organization in cattle. However, because several anatomical sample types were represented by only one or a small number of studies, study-level effects also contributed considerably to variation in community structure. Anatomical sample type remained significantly associated with bacterial community structure after accounting for study, but these findings should be interpreted in the context of study-to-study variation inherent to public 16S rRNA gene amplicon meta-analyses.

The differences in bacterial community composition observed among anatomical niches point to local physiological environments as important contributors to bacterial community assembly. Similar spatial organization has been reported across body sites in humans and other mammals, where microbial communities are shaped by niche-specific ecological conditions and host physiology [56–58]. Our results indicate that these ecological principles also apply across the bovine body, encompassing gastrointestinal, respiratory, reproductive, mammary-associated, ocular, hoof, liver, blood, and joint microbiota. The site-specific distribution of bacterial taxa, including *Prevotella* in rumen contents, *Ureaplasma* in the vagina, *Streptococcus* in the uterus, *Lactococcus* in milk, *Mycoplasma* on the ocular surface, and *Fusobacterium*, *Pasteurella*, and *Mannheimia* in their respective anatomical niches, is consistent with bacterial communities responding to the distinct physiological conditions associated with individual anatomical sites. Together, these findings support the concept that anatomical niche is an important ecological unit associated with bacterial community organization in cattle.

Anatomical structuring coexisted with substantial sharing of bacterial taxa among related anatomical sites. Despite compositional differences, related anatomical niches shared numerous bacterial taxa, particularly within reproductive and mammary-associated sites and, to a lesser extent, throughout the gastrointestinal tract. Similar patterns have been reported in host-associated microbiomes, where body sites maintain distinct microbial communities while sharing subsets of bacterial taxa through common ecological selection, shared environmental exposure, or host-associated microbial reservoirs [38, 39, 56, 57]. Together, these observations indicate that both anatomical differentiation and taxonomic sharing contribute to the organization of the bovine microbiota across anatomical niches.

The coexistence of distinct bacterial communities and shared bacterial taxa has important implications for understanding host-associated microbial ecology. Although this meta-analysis cannot determine whether shared taxa arise through microbial transmission, common ecological selection, shared environmental exposure, or other host-associated processes, these patterns provide a basis for investigating microbial reservoirs, source–sink relationships, and microbiota-mediated interactions among anatomical systems. These findings also align with the broader concept of gut–organ axes, in which microbiota can influence distant organs through microbial metabolites, immune signaling, endocrine pathways, and other host physiological responses rather than through direct microbial movement alone [33, 59–61].

### Gastrointestinal tract microbiota

The gastrointestinal tract contained multiple compartment-associated bacterial communities rather than a single homogeneous community. Bacterial community composition differed among rumen, small intestine, hindgut, and fecal samples, reflecting the distinct physiological environments associated with each gastrointestinal compartment. Similar compartment-specific organization has been reported in cattle and other ruminants, where regional physiological differences shape microbial community assembly [8, 45]. By placing gastrointestinal microbiota within a whole-body context, these results show that compartment-specific organization of the digestive tract form part of broader bacterial biogeography across the bovine body.

The distribution of available gastrointestinal datasets was highly uneven, with most samples originating from rumen contents or feces. This imbalance likely reflects the central role of the rumen in bovine digestion and nutrition, as well as the convenience and noninvasive nature of fecal sampling. In contrast, the small intestine and hindgut remain comparatively understudied, and several gastrointestinal compartments were represented by only one study in the present meta-analysis. This uneven representation limits the extent to which patterns observed for less-sampled gastrointestinal regions can be generalized and highlights the need for broader sampling throughout the digestive tract.

This sampling pattern is particularly relevant because fecal microbiota remained compositionally distinct from rumen, small intestinal, and hindgut microbiota despite sharing numerous bacterial taxa with the distal gastrointestinal tract. Because fecal samples are widely used as noninvasive surrogates for the bovine gastrointestinal microbiota, our results, together with previous work [62], indicate that fecal microbiota do not fully represent bacterial communities throughout the digestive tract. Consequently, studies investigating rumen fermentation, nutrient utilization, or enteric methane production should interpret fecal microbiome data in relation to the specific gastrointestinal compartment of interest.

Overall, gastrointestinal compartments differed in bacterial community composition and taxonomic structure and showed variation in diversity and OTU distribution. These findings suggest that understanding microbiome-mediated regulation of digestion and nutrient utilization requires consideration of the broader gastrointestinal tract rather than focusing exclusively on the rumen or fecal microbiota.

### Reproductive tract microbiota

Our results indicate that male and female reproductive microbiota are compositionally distinct while sharing many bacterial taxa across reproductive niches. Vaginal and uterine microbiota exhibited the greatest taxonomic overlap, and numerous OTUs were also shared between female and male reproductive sites. These observations suggest that male and female reproductive microbiota may be ecologically linked despite maintaining distinct bacterial communities. This pattern supports the concept that reproductive success may be influenced by the individual and interactive effects of seminal and vagino-uterine microbiota rather than by the female reproductive microbiome alone [11]. By demonstrating extensive taxonomic overlap alongside distinct bacterial communities, our results support investigating reproduction within the context of both male and female reproductive microbiota.

*Fusobacterium* provides one example of this ecological relationship. This genus was relatively abundant in seminal microbiota and was detected across multiple reproductive niches. Previous sequencing studies also identified Fusobacterium as a dominant member of healthy bull seminal microbiota [63] and detected this genus in the bovine vagina and uterus [64]. More recently, targeted qPCR and culture-based analyses confirmed that *Fusobacterium necrophorum* and *Fusobacterium varium* are prevalent members of healthy bull seminal microbiota and are also present in bovine vagino-uterine microbiota [65]. Together, these observations suggest that bacterial taxa traditionally associated with reproductive disease may also be present in healthy reproductive microbiota, with their ecological roles depending on the anatomical niche and surrounding microbial community.

The ecological relationships among reproductive microbiota may also extend beyond reproduction itself. Increasing evidence indicates that maternal and paternal microbiota can contribute to offspring microbiome establishment and developmental programming through microbial transmission, microbial metabolites, and host physiological processes [43, 66, 67]. The mechanisms underlying the observed taxonomic overlap remain unknown. Our results provide an ecological basis for investigating how maternal and paternal reproductive microbiota may contribute to offspring microbiome establishment and developmental programming.

### Respiratory tract microbiota

Our results indicate that bacterial communities are spatially organized along the bovine respiratory tract, with distinct communities associated with the nasal cavity, nasopharynx, trachea, and lungs. These observations are consistent with previous work reporting compartment-specific bacterial communities throughout the bovine respiratory tract [55]. By integrating datasets from multiple independent studies, our meta-analysis suggests that this spatial organization is a consistent feature of the bovine respiratory microbiota. Unlike previous studies focused exclusively on the respiratory tract, our meta-analysis places these compartment-specific patterns within the broader context of bacterial community organization across the bovine body.

The differences observed between the upper and lower respiratory tract likely reflect the distinct physiological environments associated with these compartments. The upper respiratory tract is continuously exposed to the external environment and supports more diverse bacterial communities, whereas the lower respiratory tract is shaped by mucociliary clearance, local immune defenses, and other host mechanisms that regulate microbial colonization [12, 58]. These physiological differences are consistent with the lower bacterial richness and diversity observed in lower respiratory tract sites compared with the nasal cavity in our meta-analysis. These observations provide an ecological basis for future studies investigating microbiome-based approaches to improve respiratory health and reduce the risk of BRD.

### Mammary gland-associated microbiota

Our results indicate that milk represents a distinct microbial niche among mammary gland-associated sites rather than simply reflecting teat-associated bacterial communities. Although teat apex, teat barrel, teat canal, and udder skin shared relatively similar bacterial communities, milk exhibited distinct bacterial community composition at both taxonomic and OTU levels. Similar compartment-specific organization has previously been reported within the bovine mammary gland [7], suggesting that milk and teat-associated niches are shaped by different physiological environments and host-associated selective factors.

The distinct microbial composition of milk may have important implications for early-life microbiome establishment. Under natural suckling conditions, newborn calves are exposed to microorganisms associated with both milk [68] and teat-associated sites [7], making the mammary gland-associated microbiota a potential source of early bacterial exposure. Increasing evidence suggests that maternal microbial exposure during early life contributes to microbiome development and immune maturation [69, 70]. These findings provide an ecological basis for investigating how colostrum processing, milk feeding practices, and milk replacer use influence early-life microbial exposure and microbiome establishment in calves.

### Other anatomical site microbiota

Compared with the gastrointestinal, respiratory, reproductive, and mammary systems, bacterial communities associated with the ocular surface, hoof, liver, blood, and joints have received considerably less attention in cattle. However, our results showed that each of these anatomical sites was associated with distinct bacterial community profiles, indicating that bacterial biogeography extends beyond the major body systems.

One important observation was the detection of bacterial DNA profiles in low-biomass tissues such as the liver, blood, and joints. The detection of microbial DNA in these anatomically protected tissues remains an area of active investigation because of their low microbial biomass and the technical challenges associated with contamination or background DNA during sample collection and sequencing. Nevertheless, bacterial DNA has been reported in healthy blood and liver in both humans and cattle [18, 19, 21], and liver-associated bacterial profiles have also been identified in healthy calves [71]. Whether these signals represent resident microbiota, transient bacterial populations, circulating bacterial DNA, or technical/background contamination remains unknown. Our results support continued investigation of these understudied anatomical niches, particularly using study designs that include appropriate negative controls and low-biomass microbiome safeguards.

Several bacterial genera were also detected across multiple anatomical sites despite the distinct bacterial community profiles associated with each niche. For example, *Moraxella* predominated in the ocular microbiota but was also detected in the respiratory tract, whereas *Fusobacterium* was abundant in liver samples and was also identified in the reproductive tract and hoof microbiota. Likewise, *Mycoplasma* occurred on the ocular surface, in the respiratory tract, and in joint samples. Similar observations have been reported in previous studies, where the ecological roles of these bacteria depend on the anatomical niche, surrounding microbial community, and host physiological state rather than the presence of a bacterial genus alone [16, 17, 22, 30, 72]. These findings highlight the importance of interpreting bacterial taxa within their ecological context rather than classifying them solely as commensals or pathogens.

### Limitations of the meta-analysis

This study has limitations inherent to public 16S rRNA gene amplicon meta-analyses. Included datasets differed in study design, geographic origin, cattle type, sample size, sequencing platform, and targeted 16S rRNA gene region, all of which can contribute to technical and biological variation in microbiota profiles [73]. In addition, several sample types were represented by only one or a small number of studies, limiting the ability to fully separate anatomical effects from study-specific effects. Although closed-reference OTU picking was used to facilitate comparisons across studies, this approach excludes sequences that do not match the reference database and limits taxonomic resolution. The analysis was also limited by the availability and consistency of public metadata, including animal age, diet, health status, management, antimicrobial exposure, sampling method, and other host or environmental factors. Finally, results from low-biomass tissues such as blood, liver, and joints should be interpreted cautiously because negative controls were not consistently available across studies and contaminant or background DNA can strongly influence sequence-based microbiota profiles from low-biomass samples [74, 75].

In summary, in this meta-analysis, we integrated publicly available 16S rRNA gene sequencing data from 5,637 samples from 47 studies to characterize bacterial biogeography across 27 anatomical niches in cattle. Our results identified patterns of anatomical structuring of bacterial communities throughout the bovine body while also identifying bacterial taxa shared among related anatomical niches, particularly within the reproductive and mammary-associated systems. Together, our study provides, to our knowledge, the first large-scale, whole-body characterization of bacterial biogeography in cattle and establishes a valuable baseline for future studies of bovine microbial ecology and host–microbiome interactions.

## Acknowledgements

We are grateful to Justine Kilama for his help creating Figure 1.

## Author Contributions

G.A. conducted the literature search, obtained and curated the sequencing datasets, performed the bioinformatic and statistical analyses, interpreted the results, and drafted the manuscript. D.B.H. contributed to the bioinformatic analyses, data interpretation, and presentation of results, and reviewed and revised the manuscript. C.R.D. contributed to the conception of the meta-analysis and reviewed and revised the manuscript. S.A. conceived and designed the study, supervised the project, interpreted the results, provided funding support, and contributed to manuscript writing and revision.

## Data Availability Statement

No new sequencing data were generated in this study. All 16S rRNA gene sequencing datasets analyzed are publicly available in the NCBI Sequence Read Archive or European Nucleotide Archive under the BioProject accession numbers listed in Supplementary Table S1

