## Supplementary figures 1-10 for "Microbial Biogeography Across the Bovine Body: A Meta-analysis of 27 Anatomical Niches"

**Supplementary figures S1-S10**

***Microbial Biogeography Across the Bovine Body: A Meta-analysis of 27 Anatomical Niches***

Godson Aryee[1], Devin B. Holman[2], Carl R. Dahlen[1] and Samat Amat[1]*

[1]*Department of Animal Sciences, North Dakota State University, Fargo, ND, USA*

[2]*Lacombe Research and Development Centre, Agriculture and Agri-Food Canada, Lacombe, AB, Canada*

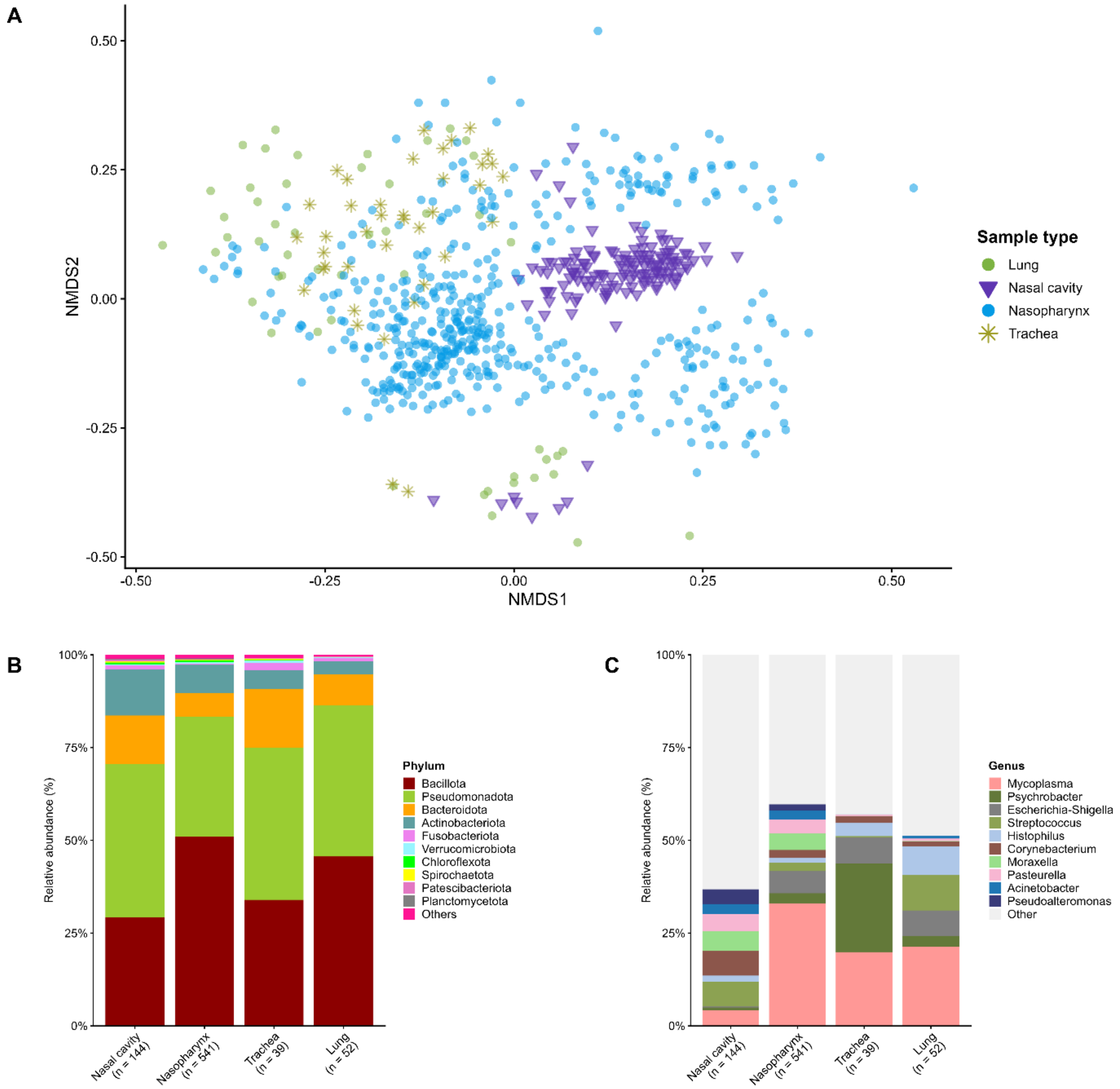


**Supplementary Figure S1.** Bacterial microbiota associated with bovine respiratory tract sample types. **A**) Non-metric multidimensional scaling (NMDS) ordination based on Bray–Curtis dissimilarities in bacterial community structure among respiratory tract samples (n = 722). **B**) Phylum-level relative abundance (%) by sample type. **C**) Genus-level relative abundance (%) by sample type. For stacked bar plots (n = 776), the most abundant taxa are shown individually and remaining taxa are grouped as “Other” or “Others.” .

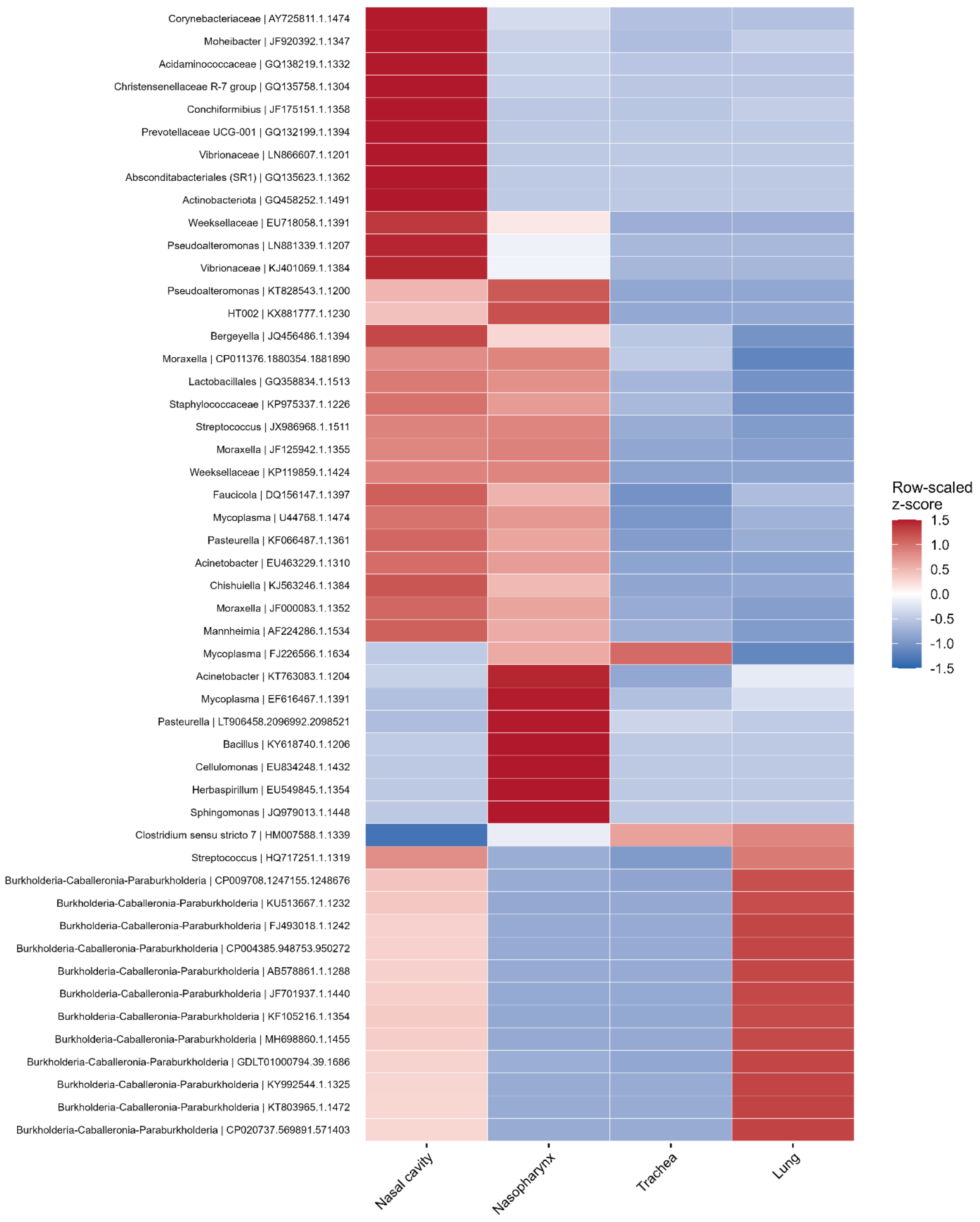


**Supplementary Figure S2.** Heatmap showing the 50 most variable operational taxonomic units (OTUs) across respiratory tract sample types. Values represent row-scaled z-scores of log10-

transformed mean relative abundance across sample types. Red indicates that an OTU had higher relative abundance in a given sample type compared with its mean abundance across respiratory tract sample types, while blue indicates lower relative abundance.

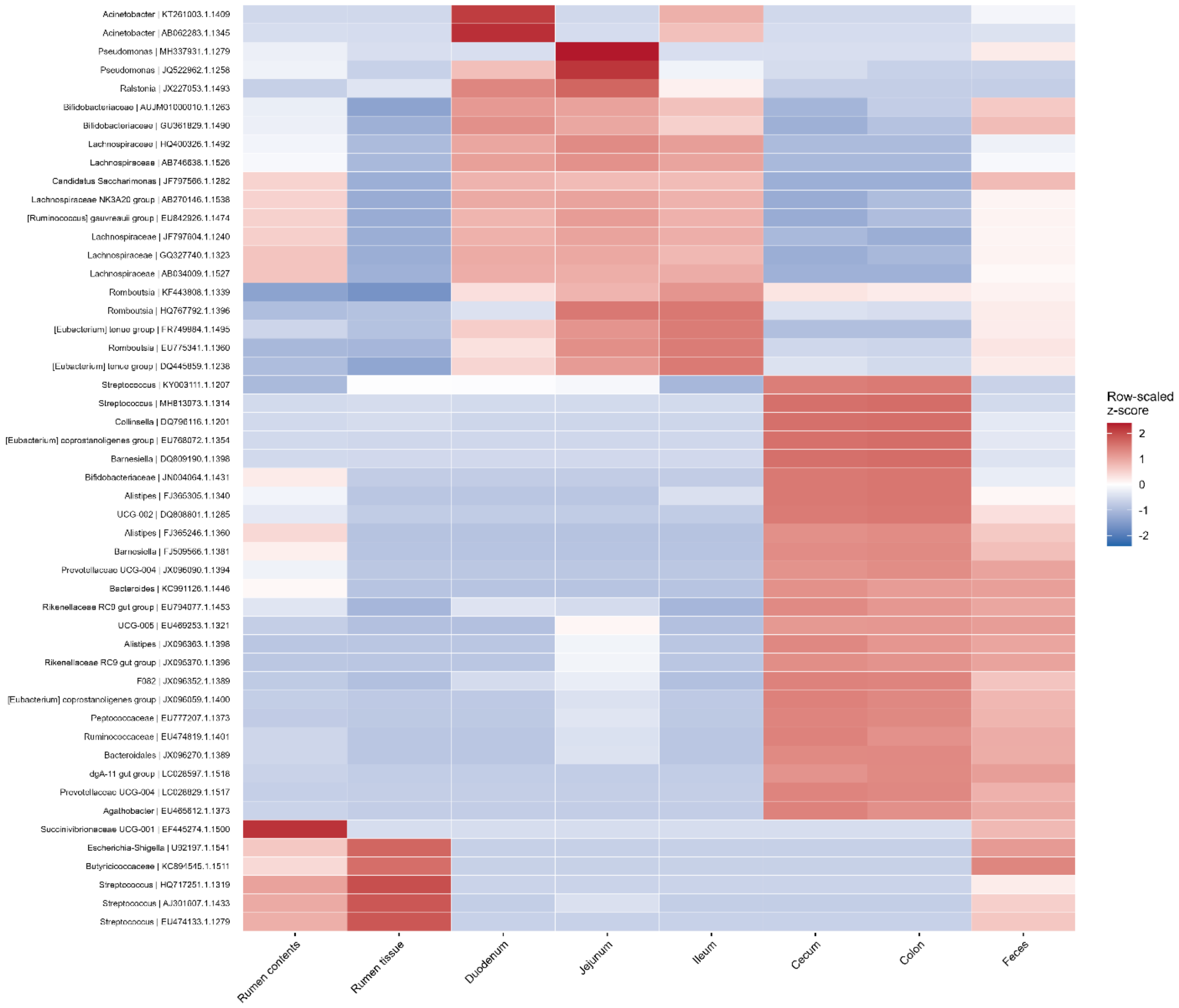


**Supplementary Figure S3.** Heatmap showing the 50 most variable operational taxonomic units (OTUs) across bovine gastroinstestinal tract sample types. Values represent row-scaled z-scores of log10-transformed mean relative abundance across sample types. Red indicates that an OTU had higher relative abundance in a given sample type compared with its mean abundance across gastrointestinal tract sample types, while blue indicates lower relative abundance.

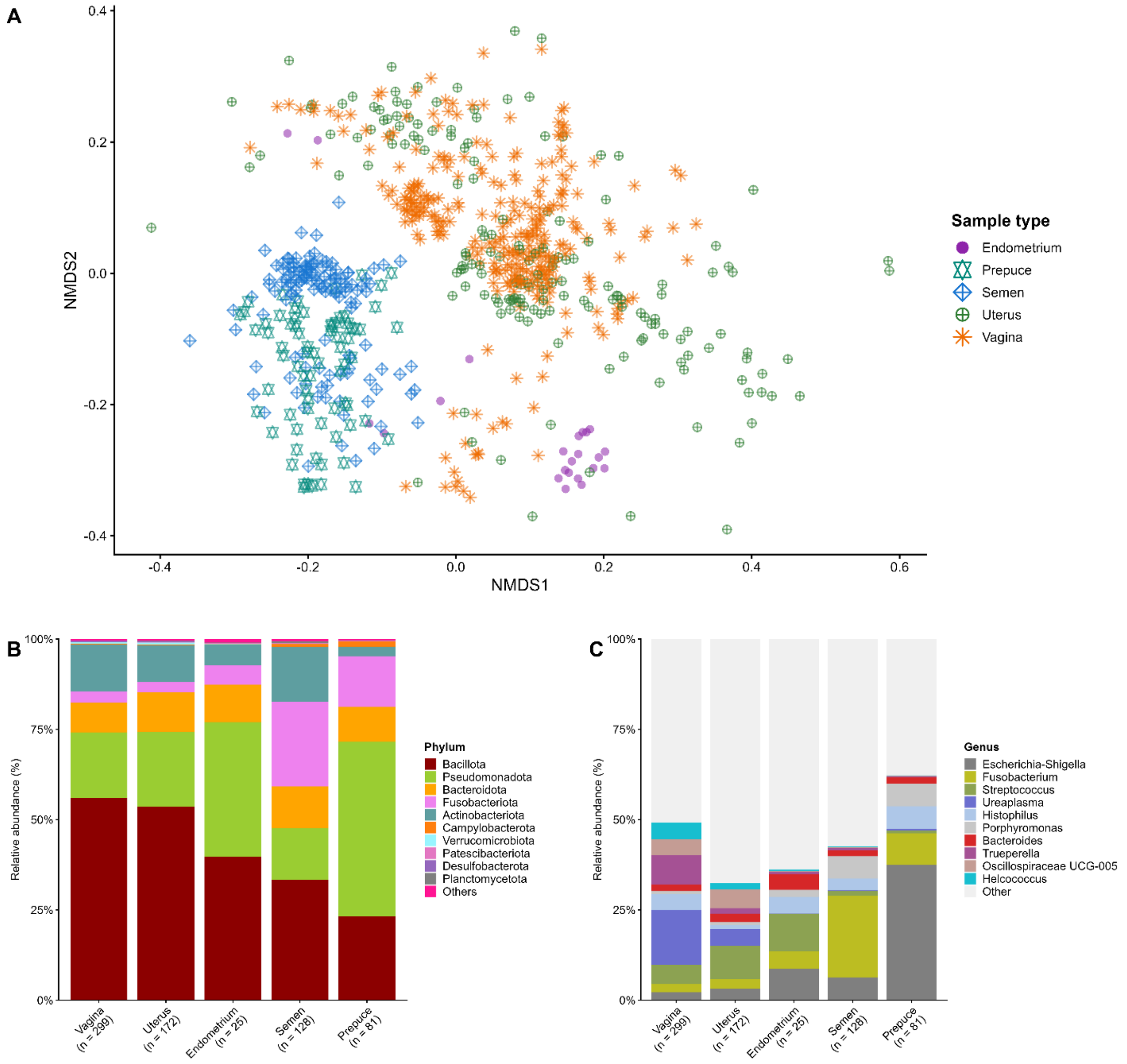


**Supplementary Figure S4.** Bacterial microbiota associated with bovine reproductive tract sample types. **A**) Non-metric multidimensional scaling (NMDS) ordination based on Bray-Curtis dissimilarities in bacterial community structure among reproductive tract samples (n = 692). **B**) Phylum-level relative abundance (%) by sample type. **C**) Genus-level relative abundance (%) by sample type. For stacked bar plots (n = 705), the most abundant taxa are shown individually and remaining taxa are grouped as “Other” or “Others.”

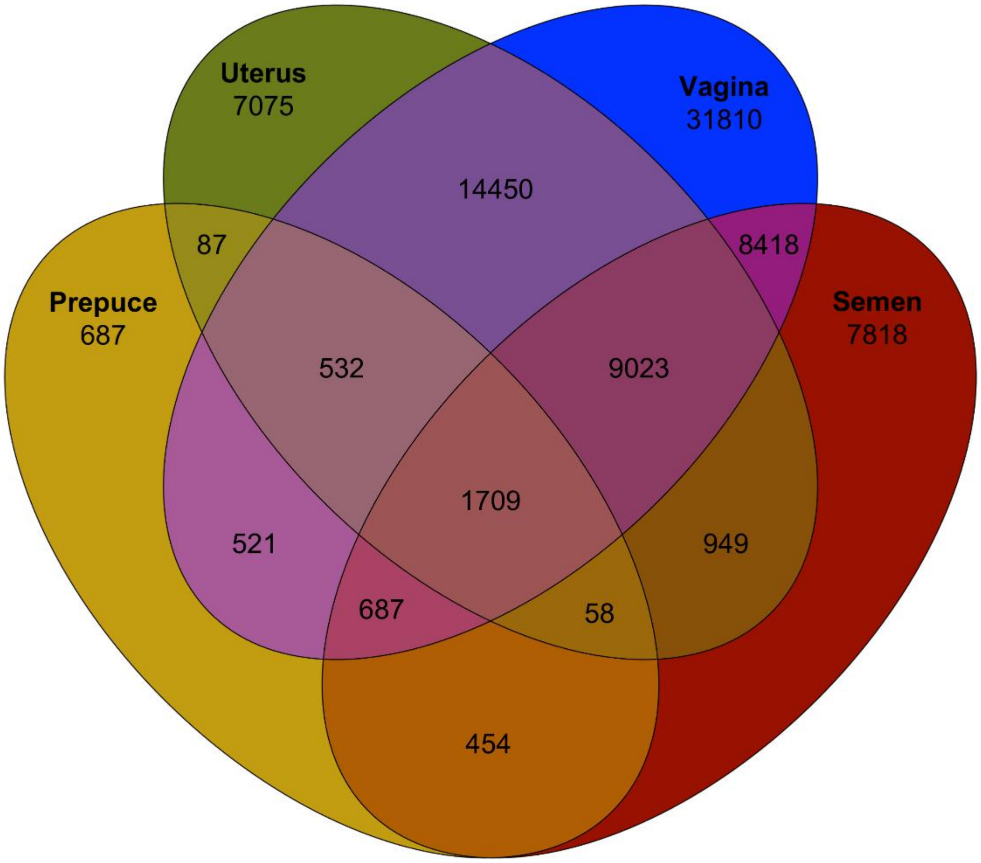


**Supplementary Figure S5.** Venn diagram showing shared and unique operational taxonomic units (OTUs) among vaginal, uterine, semen, and preputial microbiota. Numbers indicate OTUs unique to each reproductive compartment or shared among multiple compartments.

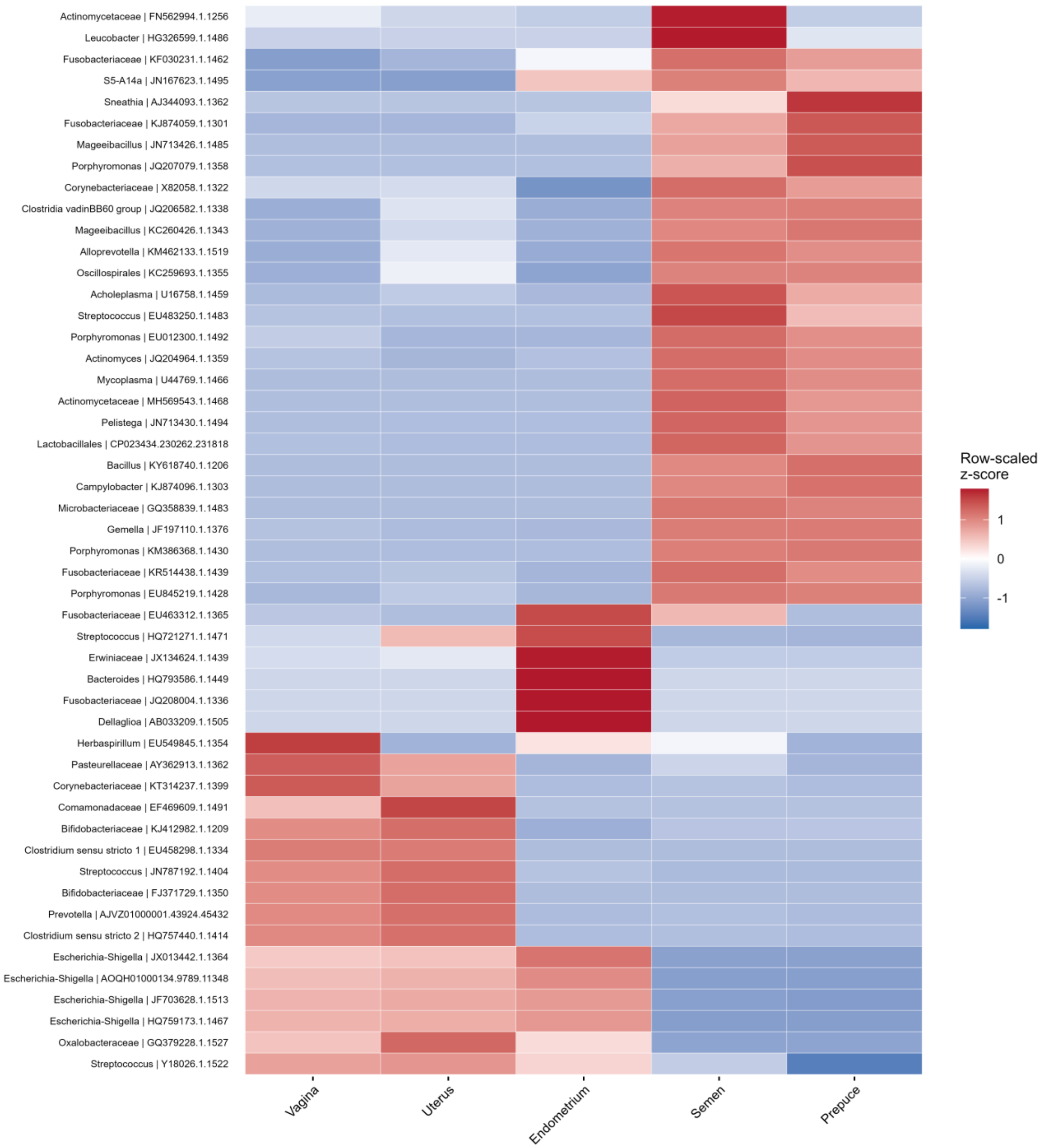


**Supplementary Figure S6.** Heatmap showing the 50 most variable operational taxonomic units (OTUs) across reproductive tract samples. Values represent row-scaled z-scores of log10-transformed mean relative abundance across sample types. Red indicates that an OTU had higher relative abundance in a given sample type compared with its mean abundance across reproductive tract sample types, while blue indicates lower relative abundance.

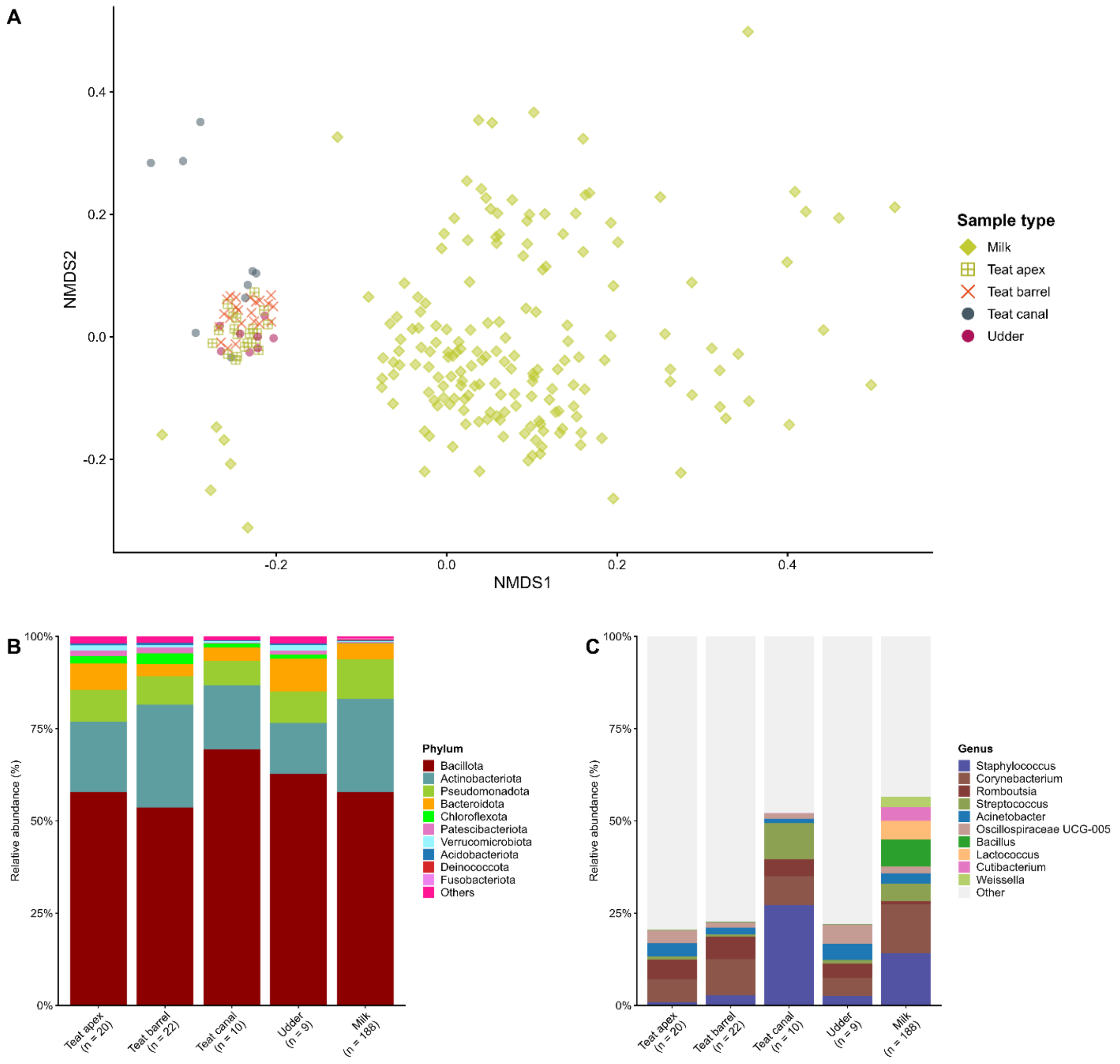


**Supplementary Figure S7**. Bacterial microbiota of mammary gland-associated sample types. **A**) Non-metric multidimensional scaling (NMDS) ordination based on Bray-Curtis dissimilarities in bacterial community structure among mammary gland-associated samples (n = 236). **B**) Phylum-level relative abundance (%) by sample type. **C**) Genus-level relative abundance (%) by sample type. For stacked bar plots (n = 249), the most abundant taxa are shown individually and remaining taxa are grouped as "Other" or "Others."

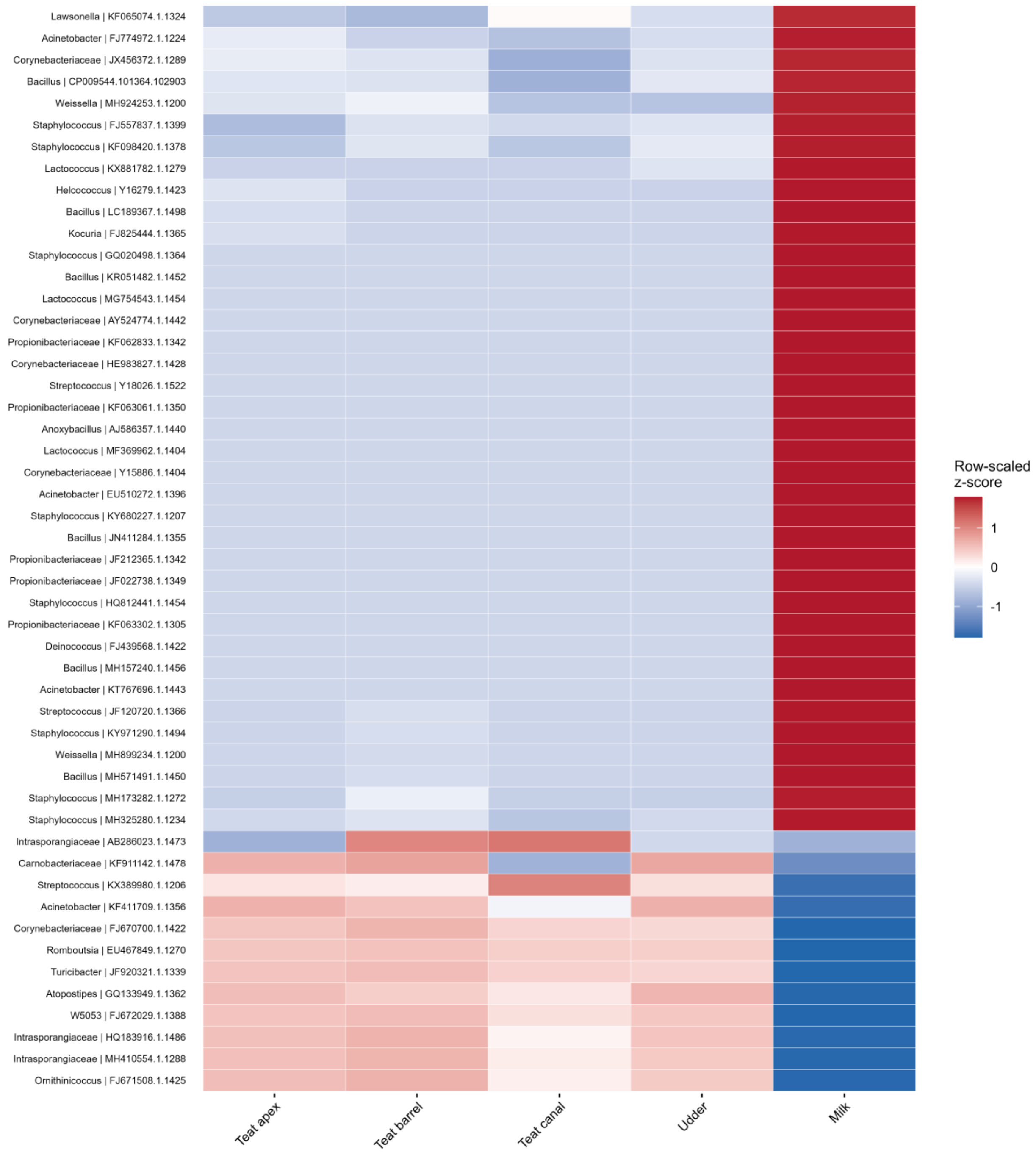


**Supplementary Figure S8.** Heatmap showing the 50 most variable operational taxonomic units (OTUs) across mammary gland-associated sample. Values represent row-scaled z-scores of log10-transformed mean relative abundance across sample types. Red indicates that an OTU had higher relative abundance in a given sample type compared with its mean abundance across mammary gland-associated sample types, while blue indicates lower relative abundance.

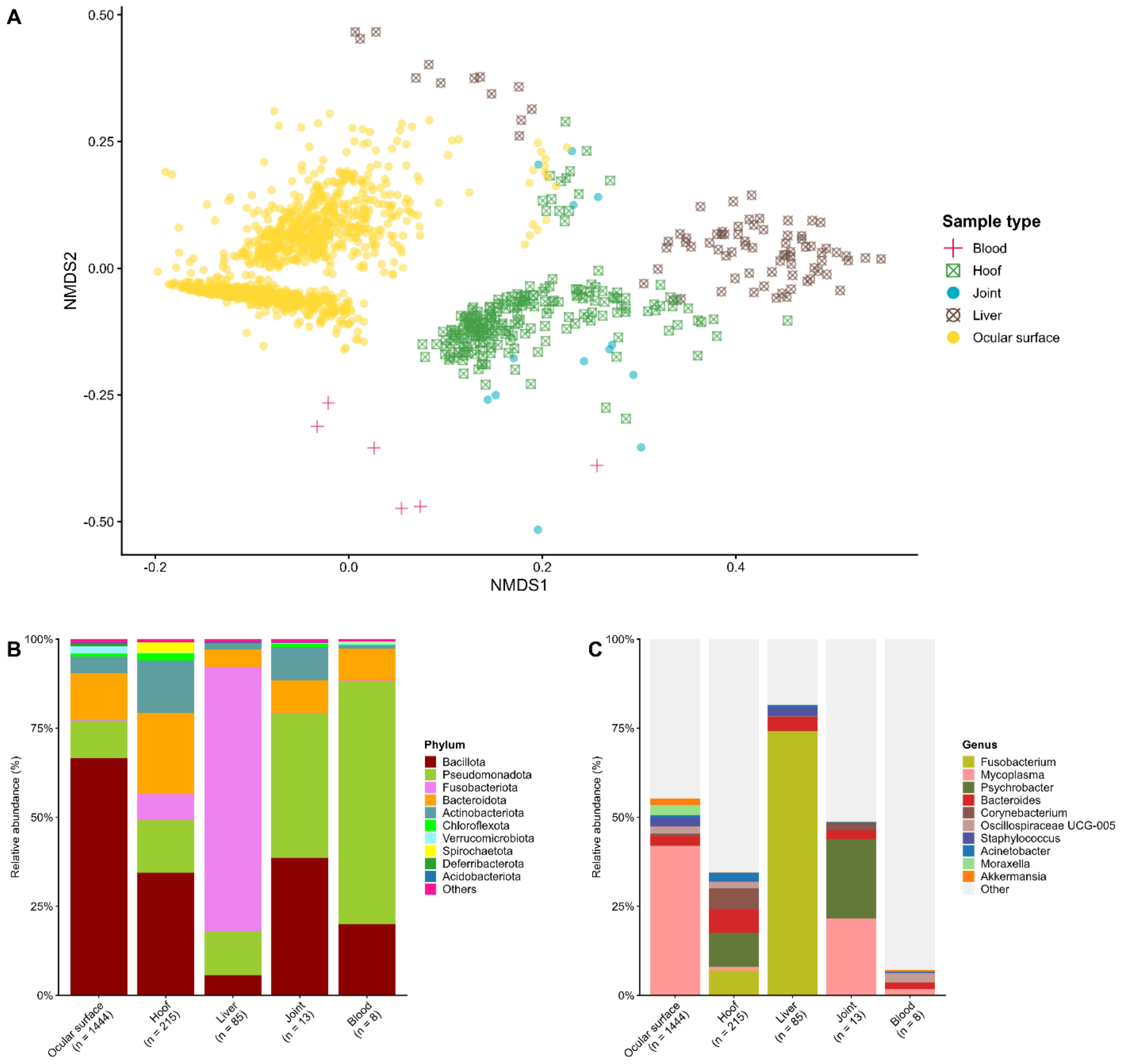


**Supplementary Figure S9**. Bacterial microbiota associated with ocular surface, hoof, liver, joint, and blood samples. **A**) Non-metric multidimensional scaling (NMDS) ordination based on Bray-Curtis dissimilarities in bacterial community structure among ocular surface, hoof, liver, joint, and blood samples (n = 1,525). **B**) Phylum-level relative abundance (%) by sample type. **C**) Genus-level relative abundance (%) by sample type. For stacked bar plots (n = 1,765), the most abundant taxa are shown individually and remaining taxa are grouped as “Other” or “Others.”

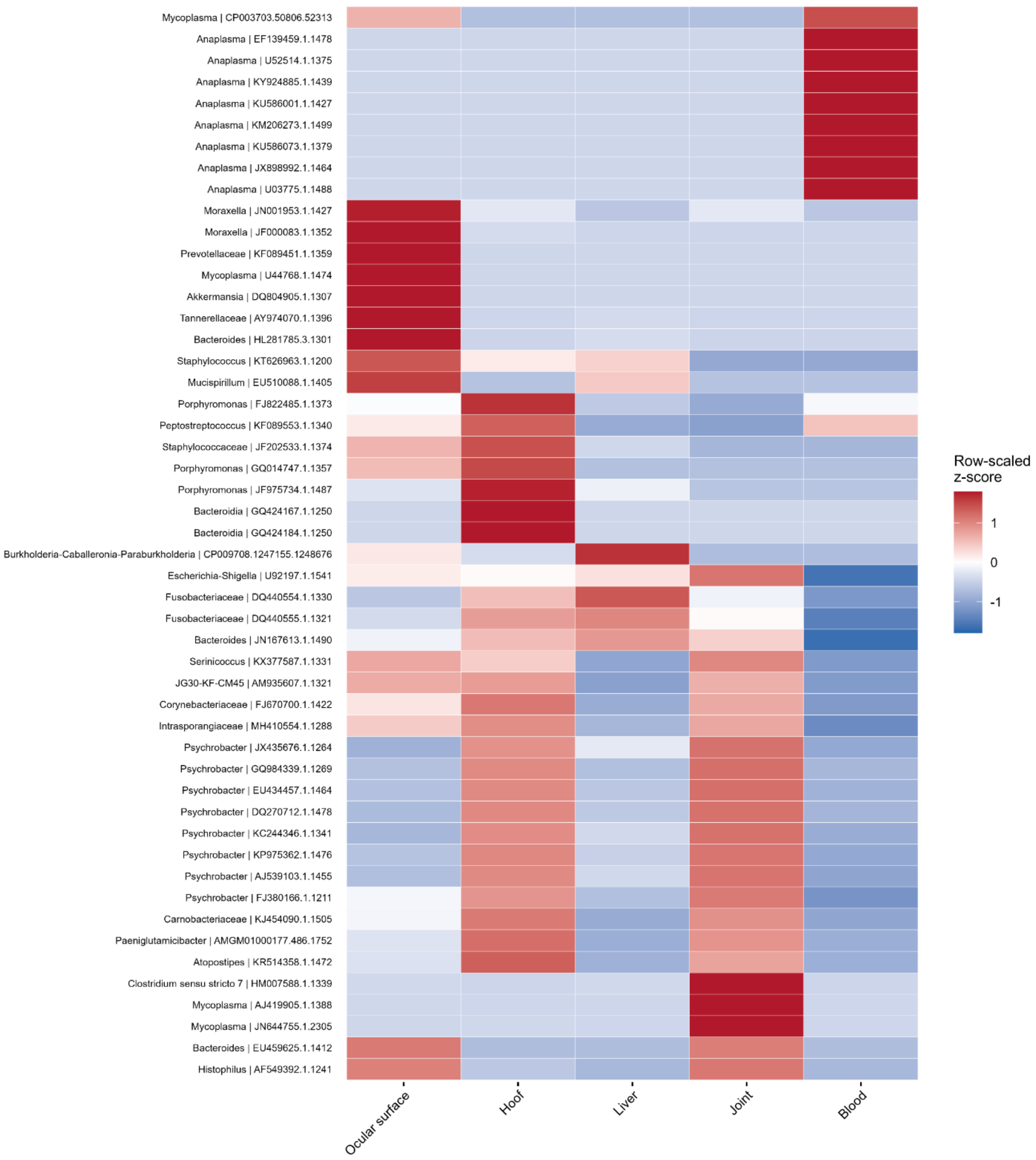


**Supplementary Figure S10**. Heatmap showing the 50 most variable operational taxonomic units (OTUs) across ocular surface, hoof, liver, joint, and blood samples. Values represent row-scaled z-scores of log10-transformed mean relative abundance across sample types. Red indicates that an

OTU had higher relative abundance in a given sample type compared with its mean abundance across sample types, while blue indicates lower relative abundance.
